# Comparative transcriptomics reveals human-specific cortical features

**DOI:** 10.1101/2022.09.19.508480

**Authors:** Nikolas L. Jorstad, Janet H.T. Song, David Exposito-Alonso, Hamsini Suresh, Nathan Castro, Fenna M. Krienen, Anna Marie Yanny, Jennie Close, Emily Gelfand, Kyle J. Travaglini, Soumyadeep Basu, Marc Beaudin, Darren Bertagnolli, Megan Crow, Song-Lin Ding, Jeroen Eggermont, Alexandra Glandon, Jeff Goldy, Thomas Kroes, Brian Long, Delissa McMillen, Trangthanh Pham, Christine Rimorin, Kimberly Siletti, Saroja Somasundaram, Michael Tieu, Amy Torkelson, Katelyn Ward, Guoping Feng, William D. Hopkins, Thomas Höllt, C. Dirk Keene, Sten Linnarsson, Steven A. McCarroll, Boudewijn P. Lelieveldt, Chet C. Sherwood, Kimberly Smith, Christopher A. Walsh, Alexander Dobin, Jesse Gillis, Ed S. Lein, Rebecca D. Hodge, Trygve E. Bakken

**Affiliations:** Allen Institute for Brain Science; Seattle, WA, 98109; Division of Genetics and Genomics and HHMI, Boston Children’s Hospital, Harvard Medical School; Boston, MA; Cold Spring Harbor Laboratory; Cold Spring Harbor, NY; Department of Genetics, Harvard Medical School; 77 Avenue Louis Pasteur Boston MA 02115; LKEB, Dept of Radiology, Leiden University Medical Center; Leiden, The Netherlands; Computer Graphics and Visualization Group, Delft University of Technology; Delft, The Netherlands; Stanley Institute for Cognitive Genomics, Cold Spring Harbor Laboratory; Cold Spring Harbor, NY; Department of Medical Biochemistry and Biophysics, Karolinska Institutet; Stockholm, Sweden; McGovern Institute for Brain Research, Massachusetts Institute of Technology; Cambridge, Massachusetts 02139; Department of Brain and Cognitive Sciences, Massachusetts Institute of Technology; Cambridge, Massachusetts 02139; Stanley Center for Psychiatric Research, Broad Institute of MIT and Harvard; Cambridge, Massachusetts 02142; Keeling Center for Comparative Medicine and Research, University of Texas, MD Anderson Cancer Center; Houston, TX; Department of Laboratory Medicine and Pathology, University of Washington; Seattle, WA, USA; Broad Institute of MIT and Harvard; Cambridge, MA 02142; Pattern Recognition and Bioinformatics group, Delft University of Technology; Delft, The Netherlands; Department of Anthropology, The George Washington University; Washington, DC; Department of Physiology, University of Toronto; Toronto, Canada

## Abstract

Humans have unique cognitive abilities among primates, including language, but their molecular, cellular, and circuit substrates are poorly understood. We used comparative single nucleus transcriptomics in adult humans, chimpanzees, gorillas, rhesus macaques, and common marmosets from the middle temporal gyrus (MTG) to understand human-specific features of cellular and molecular organization. Human, chimpanzee, and gorilla MTG showed highly similar cell type composition and laminar organization, and a large shift in proportions of deep layer intratelencephalic-projecting neurons compared to macaque and marmoset. Species differences in gene expression generally mirrored evolutionary distance and were seen in all cell types, although chimpanzees were more similar to gorillas than humans, consistent with faster divergence along the human lineage. Microglia, astrocytes, and oligodendrocytes showed accelerated gene expression changes compared to neurons or oligodendrocyte precursor cells, indicating either relaxed evolutionary constraints or positive selection in these cell types. Only a few hundred genes showed human-specific patterning in all or specific cell types, and were significantly enriched near human accelerated regions (HARs) and conserved deletions (hCONDELS) and in cell adhesion and intercellular signaling pathways. These results suggest that relatively few cellular and molecular changes uniquely define adult human cortical structure, particularly by affecting circuit connectivity and glial cell function.

## Main Text

Humans have unique cognitive abilities among primates, including our closest evolutionary cousins, chimpanzees and other great apes. For example, humans have the capacity for vocal learning that requires a highly interconnected set of brain regions, including the middle temporal gyrus (MTG) region of the neocortex that integrates multimodal sensory information and is critical for visual and auditory language comprehension (*1, 2*). Human MTG is larger and more connected to other language-associated cortical areas compared to chimpanzees and other non-human primates (NHPs) (*3–5*). These gross anatomical changes may be accompanied by changes in the molecular programs of cortical neurons and non-neuronal cells. Indeed, previous work has identified hundreds of genes with up- or down-regulated expression in the cortex of humans compared to chimpanzees and other primates (*6–9*) but have been limited to comparing broad populations of cells or have lacked a second great ape species to study changes specific to the human lineage.

Single nucleus RNA-sequencing has enabled generation of high-resolution transcriptomic taxonomies of cell types in neocortex and other brain regions. Comparative analyses have established homologous cell types across mammals, including human and NHPs, and identified conserved and specialized features: cellular proportions (*10*), spatial distributions (*11*), and transcriptomic and epigenomic profiles (*12*). In this study, we profiled over 570,000 single nuclei using RNA-sequencing from MTG of 5 species: human, two great apes (chimpanzee and gorilla), a cercopithecid monkey (rhesus macaque), and a platyrrhine monkey (common marmoset). This represents approximately 40 million years of evolution since these primate species shared a last common ancestor, and encompasses the relatively recent divergence of the human lineage from that of chimpanzees at 6 million years ago.

We defined cell type taxonomies for each species and a consensus taxonomy of 57 homologous cell types that were conserved across these haplorhine primates. This enabled comparison of the cellular architecture of cortex in humans to a representative sample of non-human primates at unprecedented resolution to disentangle evolutionary changes in cellular composition from gene expression profiles. Including gorillas, whose ancestry branched from that leading to humans and other great apes approximately 9 million years ago, enabled testing for faster evolution on the human lineage and inference of differences between human and chimpanzee that are derived (novel) in humans. Including two phylogenetically diverse monkey species enabled identification of cellular specializations that humans share with other great apes that may contribute to our enhanced cognitive abilities. Finally, establishing putative links between HARs and hCONDELs and human expression specializations by leveraging recently generated datasets of the *in vitro* activity of HARs (*13*) and cell type-specific chromatin folding (*14, 15*) identifies a subset of changes that may be adaptive.

### Within-species cell type taxonomies

MTG cortical samples were collected from post-mortem adult male and female human, chimpanzee (*Pan troglodytes*), gorilla (*Gorilla gorilla*), rhesus macaque (*Macaca mulatta*), and common marmoset (*Callithrix jacchus*) individuals for single nucleus RNA-sequencing (**Fig. 1B**). MTG was identified in each species using gross anatomical landmarks. Layer dissections for human, chimpanzee, and gorilla datasets were identified and sampled as previously described (*16*). MTG slabs were sectioned, stained with fluorescent Nissl, and layers were microdissected and processed separately for nuclear isolation.

**Fig. 1.**
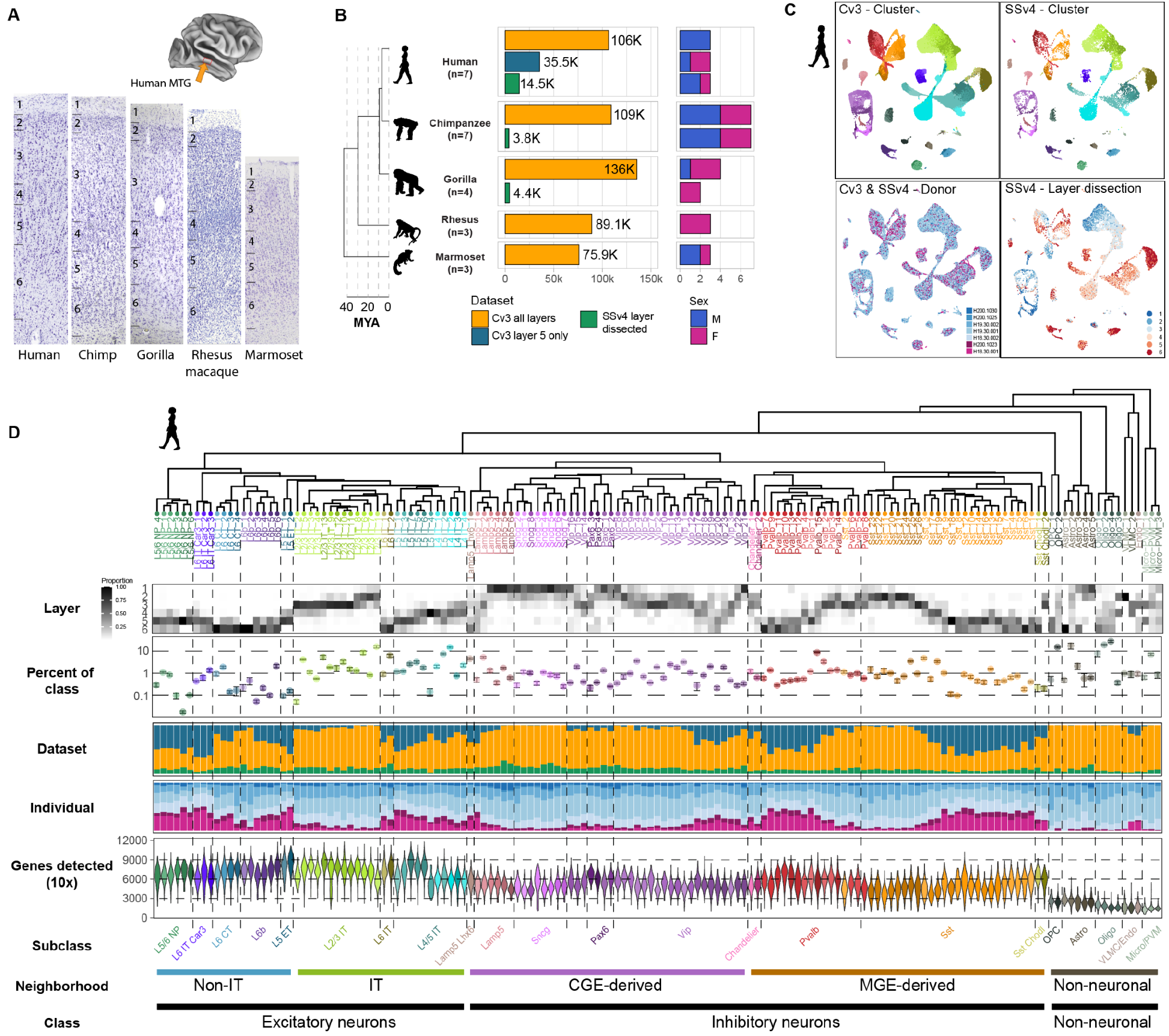
Transcriptomic cell type taxonomies of human and NHP MTG. **(A)** Approximate MTG region dissected from human brain (inset). Representative Nissl-stained cross-sections of MTG in the five species profiled. **(B)** Phylogeny of species (left; MYA, millions of years ago) and barplots of QC-passing nuclei (center) and sampled individuals (right) for each dataset. **(C)** UMAP plots of single nuclei from human MTG integrated across individuals and RNA-seq technologies and colored by cluster, individual id, and laminar dissections. **(D)** Dendrogram of human taxonomy from Cv3 cluster median expression using top 3000 high-variance genes, top. Heatmap of relative laminar distributions of cell types from SSv4 layer dissections. Dot plot of cell type abundance represented as a proportion of class (excitatory, inhibitory, glia). Error bars denote standard deviation across Cv3 individuals (L5 only dissection excluded). Barplots indicate the proportion of each cluster that is composed of Cv3 all layers, Cv3 layer 5 only, and SSv4 layer dissected datasets. Barplots indicating the proportion of each cluster that is composed of each individual. Violin plots showing the number of unique genes detected from Cv3 datasets.

For humans, single nuclei from 7 individuals contributed to three RNA-seq datasets: a Chromium 10x v3 (Cv3) dataset sampled from all 6 cortical layers (n = 107k nuclei); a Cv3 dataset sampled from micro-dissected layer 5 to capture rare excitatory neuron types (n = 36k); and our previously characterized (*16*) SMARTseq v4 (SSv4) dataset of six micro-dissected layers (n = 14.5k). Chimpanzee (n = 7 individuals) datasets included Cv3 across layers (n = 109k nuclei) and SSv4 layer dissections (n = 3.9k), and gorilla (n = 4) datasets included Cv3 (n = 136k) and SSv4 (n = 4.4k). Macaque (n = 3) and marmoset (n = 3) datasets included Cv3 from all layers (n = 89.7k and 76.9k nuclei, respectively). All nuclei preparations were stained for the pan-neuronal marker NeuN and FACS-purified to enrich for neurons over non-neuronal cells. Samples containing 90% NeuN+ (neurons) and 10% NeuN-(non-neuronal cells) nuclei were used for library preparations and sequencing. Nuclei from Cv3 experiments were sequenced to a saturation target of 60%, resulting in approximately 120k reads per nucleus. Nuclei from SSv4 experiments were sequenced to a target of 500k reads per nucleus.

Each species was independently analyzed to generate a ‘within-species’ taxonomy of cell types. First, datasets were annotated with cell subclass labels from our recently described human primary motor cortex (M1) taxonomy (*12*) using Seurat (*17*). Cell types were grouped into five neighborhoods – intratelencephalic (IT)-projecting and non-IT-projecting excitatory neurons, CGE- and MGE-derived interneurons, and non-neuronal cells – that were analyzed separately. High-quality nuclei were normalized using SCTransform (*18*) and integrated across individuals and data modalities using canonical correlation analysis. Human nuclei were well-mixed across the three datasets and across individuals (**Fig. 1C**), and similar mixing was observed for the other species (**Figs. S1, S2**). The integrated space was clustered into small ‘metacells’ that were merged into 151 clusters (**Fig. 1D**) that included nuclei from all datasets and individuals. Cell types had robust gene detection (neuronal, median 3000 to 9000 genes; non-neuronal, 1500 to 3000) and were often rare (less than 1-2% of the cell class) and restricted to one or two layers (**Table S1**). Single nuclei from the other four species were clustered using identical parameters, resulting in 109 clusters in chimpanzees (**Fig. S1A**), 116 in gorillas (**Fig. S1B**), 120 in macaques (**Fig. S2A**), and 104 in marmosets (**Fig. S2B**). Significantly, human had the most cell type diversity (151 clusters), although variation could have been driven by technical factors: sampled individuals (only female macaques), tissue dissections (layer 5 enrichment for humans), RNA-seq method (SSv4 included for great apes), and genome annotation quality.

Species cell types were hierarchically organized into dendrograms based on transcriptomic similarity (**Fig. 1D, S1, S2**) and grouped into three major cell classes: excitatory (glutamatergic) neurons, inhibitory (GABAergic) neurons, and non-neuronal cells. Each of the three major classes were further divided into cell neighborhoods and subclasses based on integrated analysis of marker gene expression, layer dissections, and comparison to published cortical cell types (*12*). In total, we identified 24 conserved subclasses (18 neuronal, 6 non-neuronal) (**Fig. S4A**) that were used as a prefix for cell type labels.

Inhibitory neurons comprised five CGE-derived subclasses (Lamp5 Lhx6, Lamp5, Vip, Pax6, and Sncg) expressing the marker *ADARB2* and four MGE-derived subclasses (Chandelier, Pvalb, Sst, and Sst Chodl) expressing *LHX6*. Excitatory neurons include five intratelencephalically (IT)-projecting subclasses (L2/3 IT, L4 IT, L5 IT, L6 IT, and L6 IT Car3) and four deep layer non-IT-projecting subclasses (L5 ET, L5/6 NP, L6b, and L6 CT). Non-neuronal cells were grouped into six subclasses: astrocytes, oligodendrocyte precursor cells (OPCs), oligodendrocytes, microglia and perivascular macrophages (Micro/PVM), endothelial cells, and vascular leptomeningeal cells (VLMCs).

This human MTG taxonomy provided substantially higher cell type resolution than our previously published human cortical taxonomies (*12, 16*) due to increased sampling (155k vs. 15-85k nuclei). All cell types matched one-to-one or one-to-many, and diversity was particularly expanded for non-neuronal subclasses and several neuronal subclasses and types: L5/6 NP (6 types), L6 CT (4), L2/3 IT FREM3 (8), and SST CALB1 (9). The FREM3 transcriptomic subtypes were distributed across layers 2 and 3, consistent with reported differences in morphology and electrophysiological properties among FREM3 neurons in human MTG (*19*).

### Cell type proportion and expression changes across primates

Neuronal subclass frequencies were estimated as a proportion of excitatory and inhibitory neuron classes based on snRNA-seq sampling to account for species differences in the ratio of excitatory to inhibitory neurons (E:I ratio) (**Fig. S4B**) (*12*). Subclass proportions were highly consistent across individuals within each species and varied significantly (one-way ANOVA, P < 0.05) across species (**Fig. 2A**). Post-hoc pairwise t-tests between humans and each NHP identified up to 5-fold more L6 IT Car3, L5 ET, and *PVALB*-expressing chandelier interneurons in marmosets. Interestingly, L2/3 IT neurons had similar proportions in MTG, in contrast to the 50% expansion of L2/3 IT neurons in humans versus marmosets in M1 (*12*).

**Fig. 2.**
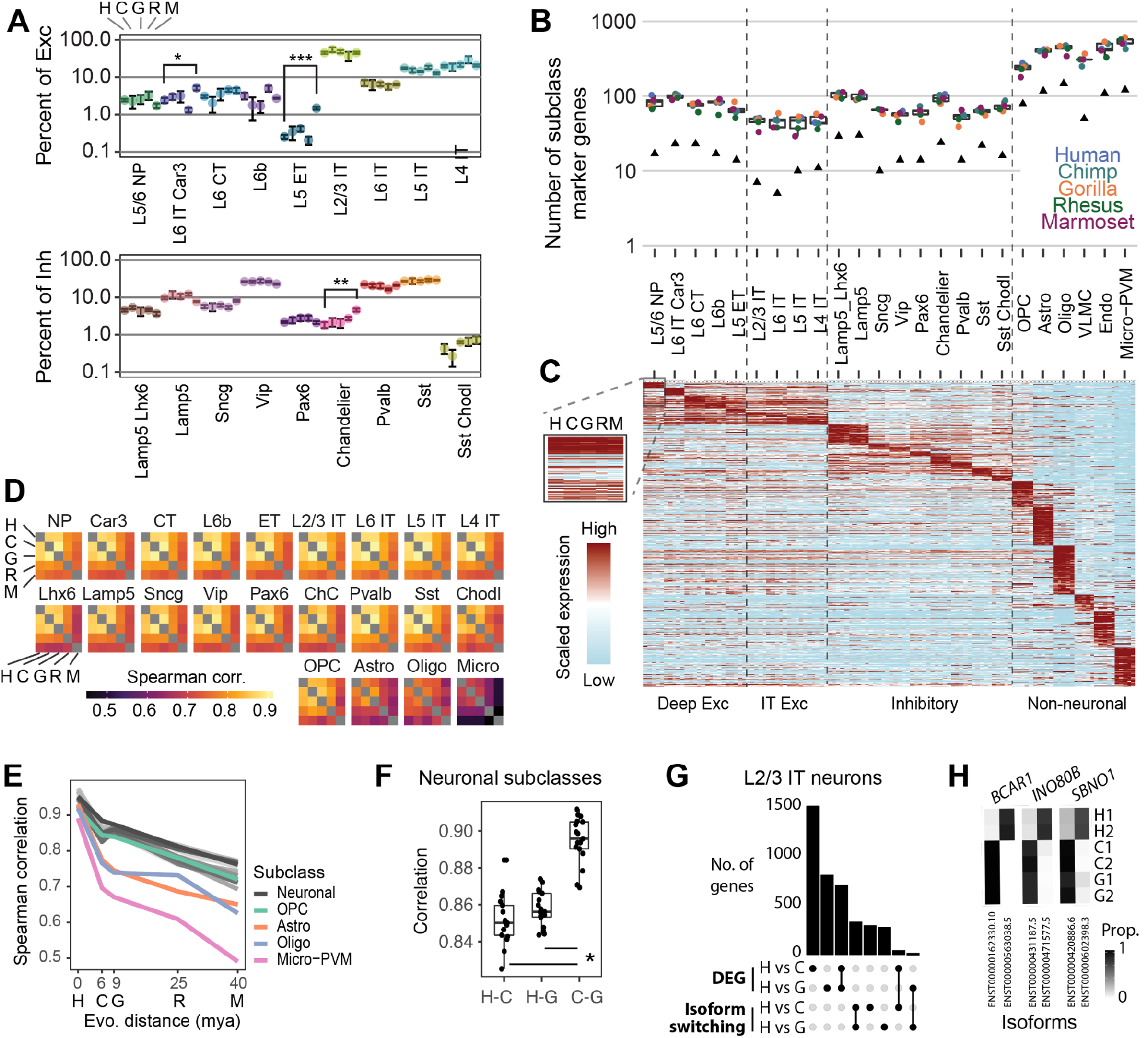
Rapid transcriptional divergence on the human lineage. **(A)** Dot plot of subclass abundance represented as a proportion of cell class. Error bars denote standard deviation across individuals. Species proportions that are significantly different from human are highlighted (two-sided t-tests; Benjamini-Hochberg corrected * P < 0.05, ** P < 0.01, *** P < 0.001). **(B)** Boxplot showing the distribution of subclass marker genes across species. Points denote the number of species markers and black triangles denote the number of conserved markers shared by all 5 species. DE genes used were significantly enriched by Wilcoxon sum rank test (adjusted p-value < 0.01) and had average log2 fold change expression greater than 1 compared to background (all other nuclei from that species). **(C)** Expression heatmap for 924 conserved markers. Expression is row-scaled for each subclass for each species. Conserved markers met criteria from **B** in all five species. **(D)** Heatmaps showing Spearman correlations of subclass median expression between species. **(E)** Line graph of subclass correlations from **D** for humans compared to each NHP as a function of the evolutionary distance of the most recent common ancestor. The zero point denotes the median correlation between human individuals for each subclass. **(F)** Boxplots of great ape pairwise correlations for neuronal subclasses from D. **(G)** Upset plot of intersections of DEGs and genes with differential isoform usage in L2/3 IT neurons between humans and chimpanzees and humans and gorillas. Differentially expressed genes were determined using a pseudo-bulk approach with DESeq2 and the Wald Test. Significantly enriched genes (adjusted p-value < 0.01) with greater than 0.5 log fold change expression in either species comparison were included. **(H)** Heatmap showing isoform proportions expressed in L2/3 IT neurons from two individuals per species for three genes (*BCAR1, INO80B*, and *SBNO1*) with human-specific switches in the main isoform.

Next, we compared the transcriptomic similarity of subclasses across primates. For each species, gene markers were defined that could reliably predict the subclass identities of cells and were filtered to include one-to-one orthologs (**Table S2**). Non-neuronal subclasses expressed hundreds of markers and were more distinct than neuronal subclasses that had 50-100 markers. Strikingly, each subclass had a similar number of markers in all species (**Fig. 2B**), but only 10-20% had strongly conserved specificity (**Fig. 2B,C**). To compare the global expression profile of subclasses across primates, we correlated normalized median expression of variable genes between each species pair for each cell subclass (excluding endothelial cells and VLMCs due to limited sampling) (**Fig. 2D**). Similarities between species reflected phylogenetic relationships, and great apes were more similar to each other than to macaque or marmoset. Marmoset was the most distantly related species and had comparably low correlations across the other NHPs and humans. Surprisingly, glial cells (except OPCs) had greater expression changes between species compared to neurons. Expression similarity decreased with evolutionary distance between human and NHPs at a similar rate across neuronal subclasses and OPCs and faster in oligodendrocytes, astrocytes, and particularly microglia (**Fig. 2E**). Some of this apparently increased evolutionary divergence may have been driven by increased variability within species, although all cell subclasses were highly similar across human individuals (median expression correlation > 0.88) (**Fig. 2E**). Indeed, glial cells (except OPCs) had more variable expression than neurons within species, but microglia and astrocytes remained significantly more divergent than neurons after normalizing for within-species variation (**Fig. S4C**). Surprisingly, chimpanzee neuronal subclasses were significantly more similar to gorilla than to human (**Fig. 2F**), despite a more recent common ancestor with human (6 versus 9 million years ago). This suggested that there was faster evolution of neurons on the human lineage since the divergence with chimpanzees. In contrast, humans, chimpanzees, and gorillas were equally divergent from macaques and marmosets (**Fig. S4D**), and there was no evidence for faster divergence on the lineage leading to great apes.

In addition to expression changes, there may be evolutionary changes in gene isoform usage. We quantified isoform expression using full-length transcript information from SSv4 RNA-seq data acquired from great apes. For each cell subclass, we identified differentially expressed genes (DEGs; **Table S3**) and genes with at least moderately high expression that strongly switched isoform usage between each pair of species (**Table S4**). Notably, there was little overlap between genes with differential expression and isoform usage for L2/3 IT neurons, the most abundant cell subclass (**Fig. 2G**). Genes with a human-specialized switch in isoform expression included *BCAR1, INO80B*, and *SBNO1* (**Fig. 2H**). *BCAR1* is a scaffold protein that is a component of the netrin signaling pathway and involved in axon guidance (*20*). *INO80B* (*21*) and *SBNO1* are involved in chromatin remodeling, and *SBNO1* contributes to brain axis development in zebrafish (*22*) and is a risk gene for intellectual disability (*23*). Interestingly, the predominant isoform of *INO80B* in human L2/3 IT neurons includes a retained intron (**Fig. S4E**) that may suppress transcription of this gene (*24*) and contribute to human specializations.

Finally, we quantified the conservation of gene expression patterns across cell types between human and NHPs. As expected, expression differences increased with evolutionary distance (**Fig. S4F**), and 75% of genes were conserved in all species (r > 0.9 in great apes; r > 0.65 in marmoset). 651 genes had highly divergent expression (r < 0.25), often in only a single species (**Fig. S4G**), such as *FAM177B* that was exclusively expressed in human microglia (**Fig. S4H**). Interestingly, a few genes had fixed derived expression in the great ape lineage. For instance, *MEPE* is a secreted calcium-binding phosphoprotein that was restricted to *PVALB*-expressing interneurons in great apes (**Fig. S4H**), and prolactin receptor (*PRLR*) had enriched expression in *SST*-expressing interneurons and L6 IT Car neurons in great apes as compared to CGE-derived interneurons in macaque and marmoset, potentially altering hormonal modulation of these neurons.

### Human specializations of glial cells

Since glial cells exhibited the most divergent gene expression changes across species (**Fig. 2D,E**), we next aimed to uncover their specialized transcriptional programs in humans compared to chimpanzees and gorillas. We first examined gene expression changes in astrocytes. There were 1189 DEGs in human astrocytes compared to chimpanzee and gorilla astrocytes. In contrast, there were fewer DEGs in the analogous chimpanzee (787) or gorilla (617) astrocyte comparisons (**Fig. 3A,B; Table S3**). Strikingly, there were three times as many highly divergent (>10-fold change in expression) astrocyte genes in humans compared to chimpanzees or gorillas. Human astrocyte DEGs were significantly enriched in synaptic signaling and protein translation pathways based on enrichment analyses using Gene Ontology (GO) (**Fig. 3C**) and Synaptic GO (SynGO) (*25*) (**Fig. 3D; Fig. S5A**) databases.

**Fig. 3.**
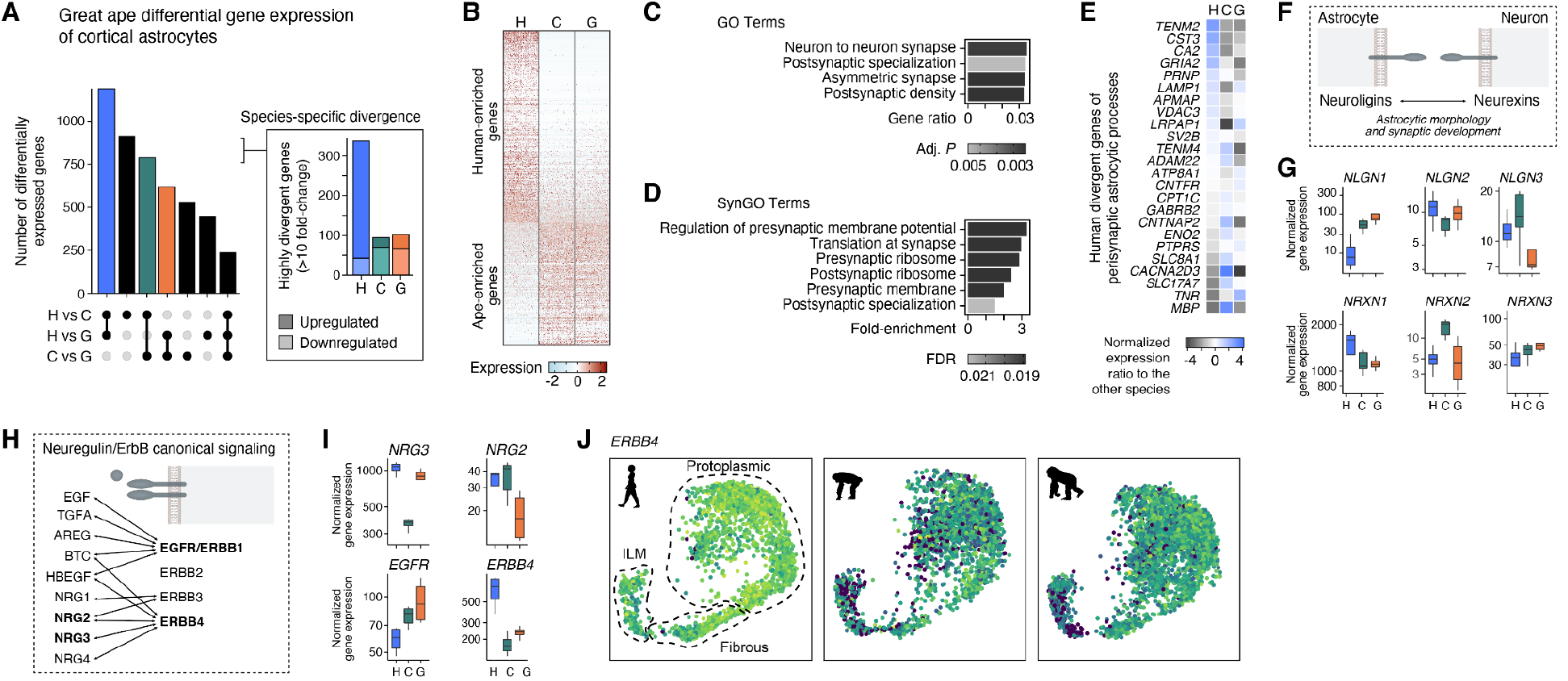
Human cortical astrocytes have specialized molecular features. **(A)** Upset plot showing the number of DEGs in cortical astrocytes for pairwise comparisons between great ape species. Inset shows the number of highly divergent genes (fold-change > 10). **(B)** Heatmap showing row-scaled expression of human versus chimpanzee and gorilla DEGs. **(C,D)** Significantly enriched GO (C) and SynGO (D) terms in the union of astrocyte DEGs from the pairwise comparison between human and chimpanzee and the pairwise comparison between human and gorilla (FDR < 0.05). **(E)** Heatmap showing human DEGs (FDR < 0.01, normalized gene count > 5) of the proteome of perisynaptic astrocytic processes (Takano et al., 2020). **(F)** Schematic illustrating the *trans*-cellular interaction of astrocytic neuroligins and neuronal neurexins that is known to play a role in astrocytic morphology and synaptic development. **(G)** Box plots showing differential gene expression of neuroligins and neurexins in astrocytes across great ape species. **(H)** Schematic illustrating ligand-receptor interactions of the neuregulin/ErbB signaling pathway. **(I)** Box plots showing differential expression of the ligands *NRG2* and *NRG3* and the receptors *EGFR* and *ERBB4* in astrocytes across great ape species. **(J)** Gene expression patterns of *ERBB4* across astrocyte subtypes and great ape species. ILM, interlaminar.

To study synapse-related astrocytic gene programs, we used a molecular database of astrocyte cell-surface molecules enriched at astrocyte-neuron junctions from an *in vivo* proteomic labeling approach in the mouse cortex (*26*). Among genes encoding 118 proteins that were robustly enriched in perisynaptic astrocytic processes, 24 genes (20%) were differentially expressed in human astrocytes compared to chimpanzee and gorilla astrocytes (**Fig. 3E**), and 47 genes (40%) had conserved expression across great apes (**Fig. S5B,C**). Interestingly, neuroligins and neurexins, ligand-receptor pairs that play a key role in astrocytic morphology and synaptic development (*27*), showed divergent expression patterns across great ape species (**Fig. 3F,G**). Other cell-adhesion gene families with well-known functions in astrocytic morphological and synaptic development also had multiple members among human astrocyte DEGs, including ephrins and their cognate receptors (*EFNA5, EPHA6*), clustered protocadherins (*PCDH9*), and teneurins (*TENM2, TENM3, TENM4*) (**Fig. S5D,E**).

In addition to cell-adhesion programs, we explored other cell-surface or secreted ligands and receptors that contribute to astrocyte function. We found that several astrocyte-secreted synaptogenic molecules such as Osteonectin (*SPARC*) and Hevin (*SPARCL1*) and ECM-related proteins (Brevican, *BCAN*; Neurocan, *NCAN*; and Phosphacan, *PTPRZ1*) were up-regulated in human astrocytes (**Fig. S5D,E**). Of note, four members of the neuregulin/ErbB signaling pathway showed differential gene expression in great ape astrocytes, with two receptors (*EGFR* and *ERBB4*) displaying expression changes in opposite directions (**Figs. 5H,I**). Interestingly, up-regulation of human *ERBB4* expression was higher in protoplasmic and fibrous astrocytes than interlaminar astrocytes (**Figs. 3J, S5F**), demonstrating that transcriptional specializations can occur in a subtype-specific fashion. Finally, glutamate AMPA receptor subunits (*GRIA1, GRIA2, GRIA4*) had more than 3-fold greater expression in human astrocytes, suggesting a human-specific responsiveness of astrocytes to glutamate (**Fig. S5G**).

We next focused our analysis on gene expression changes in microglia, which also play critical roles in cortical circuit formation (*28, 29*). Recent comparative spatial transcriptomic data indicate that microglia-neuron contacts are more prevalent in human cortical circuits compared to mice, particularly in superficial layers (*30*). We reasoned that evolutionary changes in microglial connectivity could be mediated by fine-tuning expression of cell-surface ligands and receptors. Indeed, we found that human microglia have more DEGs (328) than chimpanzee (175) or gorilla (164) microglia (**Fig. S6A,B**), and human DEGs were significantly overrepresented in GO and SynGO terms related to synaptic compartments (**Fig. S6C,D,E**). Among 58 ligand and receptor genes that were recently predicted to mediate microglia-neuron communication in the mouse cortex (*31*), we found that only 5 genes (9%) were differentially expressed in human microglia, suggesting that selective molecular processes may underlie specializations of microglia-neuron interactions in the human cortex. Remarkably, we found several disease-linked genes among human microglia DEGs (**Fig. S6F**), including up-regulation of *SNCA* (encoding alpha-synuclein) and *TMEM163* implicated in neurodegenerative disorders (*32–34*), and down-regulation of the synapse-related Rho guanine nucleotide-exchange factor Kalirin (*KALRN*) associated with neurodevelopmental and neuropsychiatric disorders (*35*) (**Fig. S6F**). We also corroborated the human-specific up-regulation of *FOXP2* and *CACNA1D* that was recently reported in the dorsolateral prefrontal cortex (*9*).

Oligodendrocytes also showed human specializations, including DEGs involved in myelin organization and cell adhesion (e.g., *CNTNAP2, LAMA2*) (**Fig. S6**). Our data confirms 145 DEGs (30% overlap) previously reported in a comparative study between human and chimpanzee oligodendrocytes and provides further insight into gene expression differences across great ape species. Notably, unlike astrocytes and microglia, human and chimpanzee oligodendrocytes had similar numbers of DEGs, although humans had more up-regulated, highly divergent DEGs (**Fig. S6G**). In summary, these findings support faster divergence of glial expression in the human lineage that parallels neuronal divergence and likely impacts interactions between glia and neurons.

### L6 IT Car3 specialization in great apes

In all species, IT-projecting excitatory neurons formed three transcriptionally distinct groups: L2/3 and L4 IT, L5 and L6 IT, and L6 IT Car3. These groups were divided into discrete cell types (**Fig. 1D, S1-2**), and related types were distributed as a gradient across cortical layers (**Fig. S7A**). The transcription factors *CUX2* and *TSHZ2*, a transcriptional repressor, were expressed in opposing gradients along this same laminar axis (**Fig. S7A,B**) with the highest *CUX2* expression in L2/3. Studies of mouse cortical development have demonstrated the importance of these TFs in determining cell type identity where *CUX2* is similarly expressed in L2/3 IT neurons and *TSHZ2* is selectively expressed in L5 NP neurons (*36*). These results in the primate cortex highlight the impact of laminar position on transcriptomic identity, and a mixture of discreteness and continuity among neuronal types that has also been reported in the adult mouse cortex (*37*).

A closer examination of L6 IT Car3 neurons revealed two distinct subtypes that expressed *CUX2* at high or low levels in the five species (**Fig. 4A**). *HTR2C* and *MGAT4C* were additional conserved markers of the High-*CUX2* subtype, and *BCL11A* and *LDB2* were markers of the Low-*CUX2* subtype (**Fig. 4B, S7C**).

**Fig. 4.**
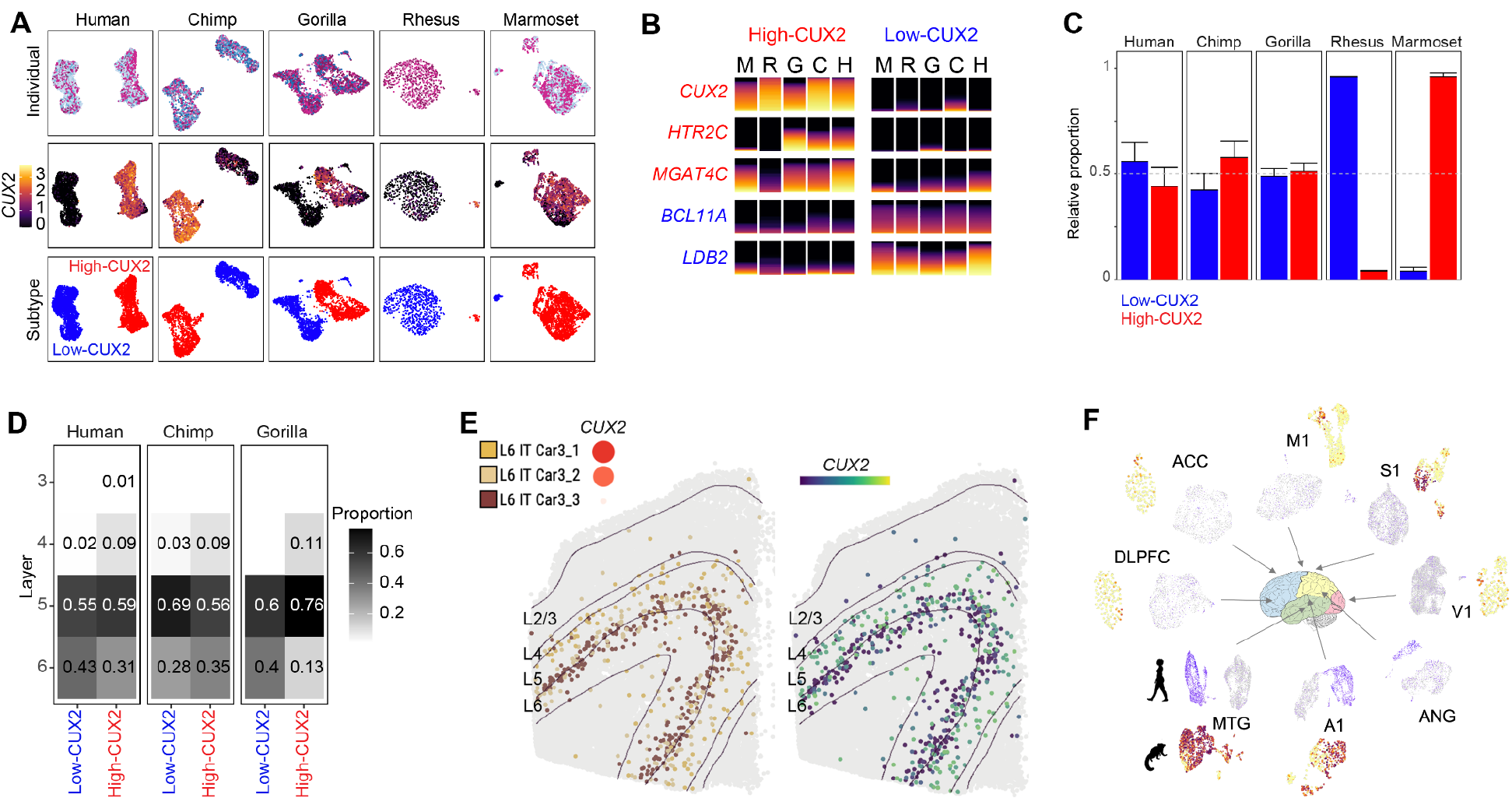
Great ape specialization of L6 IT Car3 neurons. **(A)** UMAPs of L6 IT Car neurons in each species labeled by individual, *CUX2* expression, and subtype. **(B)** Stacked bar plots of conserved marker gene expression of High-CUX2 and Low-CUX2 subtypes across the five species. **(C)** Mean (whiskers, standard deviation across individuals) proportion of two subtypes among L6 IT Car3 neurons in each species. **(D)** Heatmaps denote the proportion of nuclei dissected from each layer for each subtype across the great apes. **(E)** Neurons labeled by cell type based on expression of marker genes measured by the MERFISH spatial transcriptomics method in a human cortical slice of MTG. L6 IT Car3_1 and Car3_2 correspond to the High-CUX2 subtype, and Car3_3 corresponds to Low-CUX2 based on the average expression of *CUX2* (inset). Neurons with low expression of *CUX2* are enriched at the layer 5/6 border. **(F)** UMAPs of *CUX2* expression in L6 IT Car3 neurons from eight cortical regions in humans (purple corresponds to high expression) and seven matching regions in marmosets (dark orange corresponds to high expression). Note that the High-CUX2 subtype is more prevalent in temporal and, to a lesser extent, parietal regions.

Subtype proportions varied dramatically across the species with balanced proportions in all great ape species, mostly Low-*CUX2* in macaques, and mostly High-*CUX2* in marmosets (**Fig. 4C**). Based on laminar dissection information, Low-*CUX2* neurons were consistently enriched in deeper layers than High-*CUX2* in all three great apes (**Fig. 4D**). In human MTG, *in situ* labeling of marker genes using MERFISH (**Fig. 4E, S7D**) and RNAscope (**Fig. S7E**) validated that the Low-*CUX2* subtype was enriched at the border of layers 5 and 6, and the High-*CUX2* subtype extended from upper layer 6 to layer 4. Intriguingly, based on snRNA-seq data we collected from 7 additional regions of the human cortex (described in a companion manuscript), the Low-*CUX2* subtype was dominant in the majority of regions, and the High-*CUX2* subtype was enriched specifically in temporal cortex (MTG and primary auditory, A1) and less so in parietal cortex (angular gyrus, ANG and primary somatosensory, S1) (**Fig. 4F**). Similarly, snRNA-seq data collected from 6 additional regions of the marmoset cortex (described in a separate companion manuscript) revealed that the High-*CUX2* subtype was most enriched in temporal areas (MTG and A1) and less enriched in S1 (**Fig. 4F**).

### Consensus cell type conservation and divergence

To further investigate the canonical architecture of primate MTG, we built a transcriptomic taxonomy of high resolution consensus cell types. Starting with CGE-derived interneurons, we integrated single nucleus expression profiles across the five species based on conserved co-expression using Seurat (*17*). Within-species cell types remained distinct, and nuclei were well integrated (**Fig. 5A,B; Fig. S8**), particularly for humans and chimpanzees (**Fig. 5C**). Similar results were observed for the other cell neighborhoods (**Figs. S9-12**). Separate pairwise alignments between human and NHPs confirmed that cell type homologies were better resolved in more closely related species (**Fig. 5D**). We also found that excitatory neurons were less well integrated than inhibitory neurons, and this finding was consistent with greater species specializations of excitatory types.

**Fig. 5.**
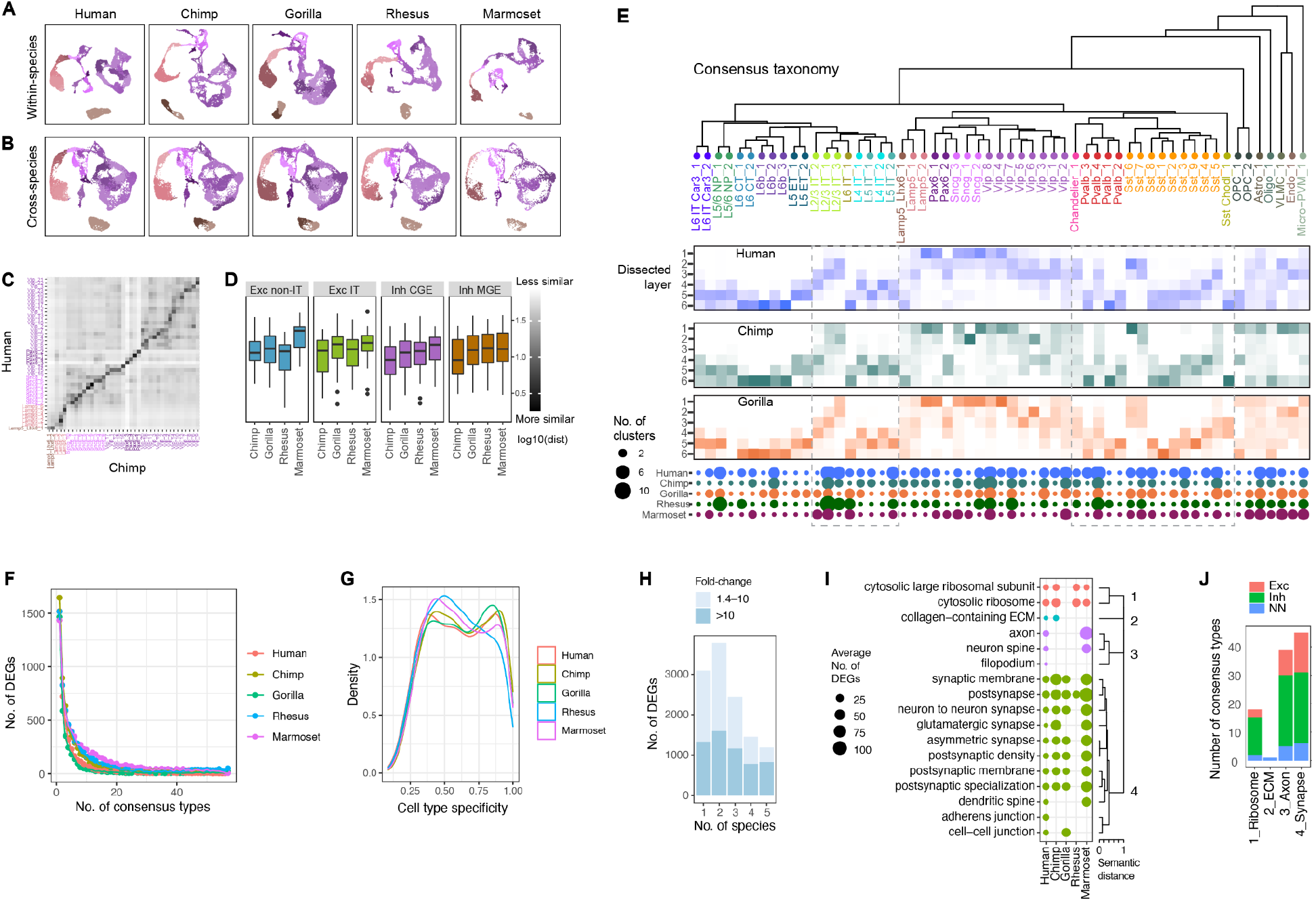
Divergent expression across conserved cell types. **(A)** UMAPs of CGE-derived interneuron expression generated for each species and colored by within-species cell types. **(B)** UMAPs of CGE interneuron expression integrated across the five species and with the same coloring as **A. (C)** Heatmap of the log-transformed Euclidean distance between human and chimpanzee cell type centroids shown in **B. (D)** Boxplots summarizing the distance to each human cluster from the nearest corresponding NHP cluster for each pairwise species integration. **(E)** Consensus taxonomy of 57 homologous cell types identified in all five species. Heatmaps denote the inferred laminar distribution of types in the great ape species based on dissections. Dot plot denotes the number of within-species clusters that are associated with each consensus type. **(F)** Distributions of the number of species DEGs that are differentially expressed across different numbers of consensus types. Most DEGs are specific to one or a few consensus types. **(G)** Cell type specificities (tau) of genes in each species that are differentially expressed in at least one type. A tau score of 0 indicates expression is evenly distributed across types, and 1 indicates a binary marker expressed in a single type. **(H)** Species DEGs including highly divergent (>10-fold change) DEGs are often differentially expressed in multiple species in different cell types. **(I)** Summary of GO enrichment analysis of species DEGs. Cellular component terms are shown that were significantly enriched for hDEGs in at least one consensus type and form four distinct groups of similar GO terms. **(J)** Summary of the number of consensus types that express hDEGs that were enriched for at least one term in the four semantic GO groups.

We established homologous cell types between all pairs of species using MetaNeighbor, a statistical framework (*38, 39*) that identified cell types that could be reliably discriminated (AUROC >0.6) from nearest neighbors in one species based on training data from the other species or that were reciprocal best matches. Pairwise cell type homologies were integrated to define 57 consensus types that included cell types identified in the five species, and a dendrogram was constructed based on transcriptomic similarities (**Fig. 5E)**. Eight consensus types represented one-to-one matches across all species, and the majority of types represented multiple matches of between two and ten within-species types. Thus, there was a conserved set of cell types in primate MTG with transcriptomic specializations of subtypes, but no distinct novel types in any species. Laminar distributions of types were remarkably conserved across the great apes, except Sst Chodl_1 was in more superficial layers of gorilla MTG and OPC_1 was present in layer 1 of chimpanzee and gorilla but not human MTG (**Fig. 5E**), although more sampling of these rare types is needed for validation.

Previous work reported the lack of transcript and protein expression of tyrosine hydroxylase (*TH*), a key enzyme in the dopamine synthesis pathway, in the neocortex of non-human African great apes including chimpanzee and gorilla (*40, 41*). Recent transcriptomic profiling of chimpanzee prefrontal cortex suggests that this represents loss of dopamine signaling in a conserved cell type rather than loss of a homologous type (*9*). In MTG, we identified 9 consensus *SST*-expressing interneuron types present in all five primates (**Fig. 5E**) that had robust sets of conserved and species-specific markers (**Fig. S13A; Table S5**). The Sst_1 consensus type was distinct from most other MGE-derived interneurons (**Fig. S13B**) and expressed *TH* in human, macaque, and marmoset neurons but not chimpanzee or gorilla neurons (**Fig. S13C-E)**. Conserved (e.g. *NCAM2, PTPRK, UNC5D*, and *CNTNAP5*) and species-specific genes were enriched in pathways for connectivity and signaling (**Fig. S13F-J**). Sst_1 was the rarest type in all primates (**Fig. S13K**) and varied from 0.3% of *SST*-expressing interneurons in gorillas and macaques to 1-3% in humans, chimpanzees, and marmosets. Interestingly, the majority of *TH*-expressing neurons belonged to different interneuron subclasses in humans (*SST*), macaques (*PVALB*), and marmosets (*VIP*) (**Fig. S13L**), and this was confirmed by *in situ* labeling of *TH*-expressing neurons in human and macaque MTG (**Fig. S13M**). Dopamine receptor expression varied across primates but did not track with predicted differences in local dopamine production (**Fig. S13N-O**).

This is likely because subcortical regions provide the majority of dopaminergic input to the neocortex and mask the effects of evolutionary changes in local input.

We tested for changes in proportions of neuronal consensus types across primates using a Bayesian model (scCODA) that accounted for the compositional nature of the data (**Fig. S14A**) (*42*). We found that the higher E:I ratio in marmosets (**Fig. S4B**) was driven by increased proportions of most excitatory types, and the lower E:I ratio in macaques was driven by increased proportions of particularly Sst and Vip interneuron types and by decreased proportions of L2/3 IT_2, L2/3 IT_3, and L6 IT Car3_2 excitatory types. There were smaller changes among the great apes, except for an increased proportion of L5/6 NP_2 neurons in humans and chimpanzees.

Next, we identified species-specialized genes by comparing consensus cell type expression for each species to all other primates. Human consensus types had a broad range (fewer than 100 to over 1000) of statistically significant DEGs (**Fig. S14B; Table S6**) that represented 1-8% of expressed genes (**Fig. S14C**). Excitatory types in deep layers (IT and non-IT) had the most human-specific DEGs (hDEGs), and non-neuronal types had the least (**Fig. S14B**). Types with fewer hDEGs had greater median fold-changes (**Fig. 14C**) that likely resulted from reduced power to detect smaller expression changes due to more variation across individuals (see **Fig. S4C**). Strikingly, many species DEGs were restricted to one or a few cell types, particularly for great apes (**Fig. 5F**). The cell type specificity of DEGs was not simply a result of expression changes in marker genes, but also selective changes in broadly expressed genes (**Fig. 5G**). hDEGs had a median 4-fold change in expression, while a few metabolism-related genes changed expression by 20-fold or more in most cell types (**Fig. S14D**). The same genes were often differentially expressed in multiple species (**Fig. 5H**) but in different cell types, and highly divergent (>10-fold) genes were usually found across all species. Species DEGs were enriched in four major pathways: ribosomal processing, extracellular matrix (ECM), axon structure, and the synapse (**Fig. 5I,J; Fig. S14E**). Ribosomal processing was primarily associated with interneurons in humans and all cell types in chimpanzees, macaques, and marmosets. Intriguingly, ECM-associated DEGs, including several laminin genes, were specific to the VLMC_1 consensus type in humans, chimpanzees, and marmosets (**Fig. S14E**) and have potential to alter the blood brain barrier as shown in a mouse model of pericyte dysfunction (*43*). Hundreds of axonal and synaptic genes were differentially expressed in most cell types in all species, and this suggests extensive molecular remodeling of connectivity and signaling during primate evolution.

### Enrichment of HARs and hCONDELs near human differentially expressed genes

Genes may change expression between species due to neutral or adaptive evolution. To investigate which hDEGs may be under positive selection, we linked hDEGs to human-specific genomic sequence changes. Because hDEGs are differentially expressed in only one or a few consensus cell types, expression changes are likely caused by sequence modifications to regulatory regions that can alter transcription in select cell types. We examined two previously identified classes of genomic regions that have changed along the human lineage: (1) human accelerated regions (HARs) that are highly conserved across mammals and have higher substitution rates in the human lineage (*13*); and (2) human conserved deletions (hCONDELs) that are highly conserved across mammals and deleted in humans (*44, 45*).

Strikingly, we find that HARs and hCONDELs are significantly (FDR < 0.05) enriched near hDEGs in virtually all consensus types (**Fig. 6A**). The proportion of hDEGs near HARs and hCONDELs is highest for non-neuronal consensus types such as VLMCs (VLMC_1), microglia (Micro-PVM_1), and oligodendrocytes (Oligo_1), likely due to the larger intronic and flanking intergenic regions of hDEGs in these cell types (**Fig. S19**).

**Fig. 6.**
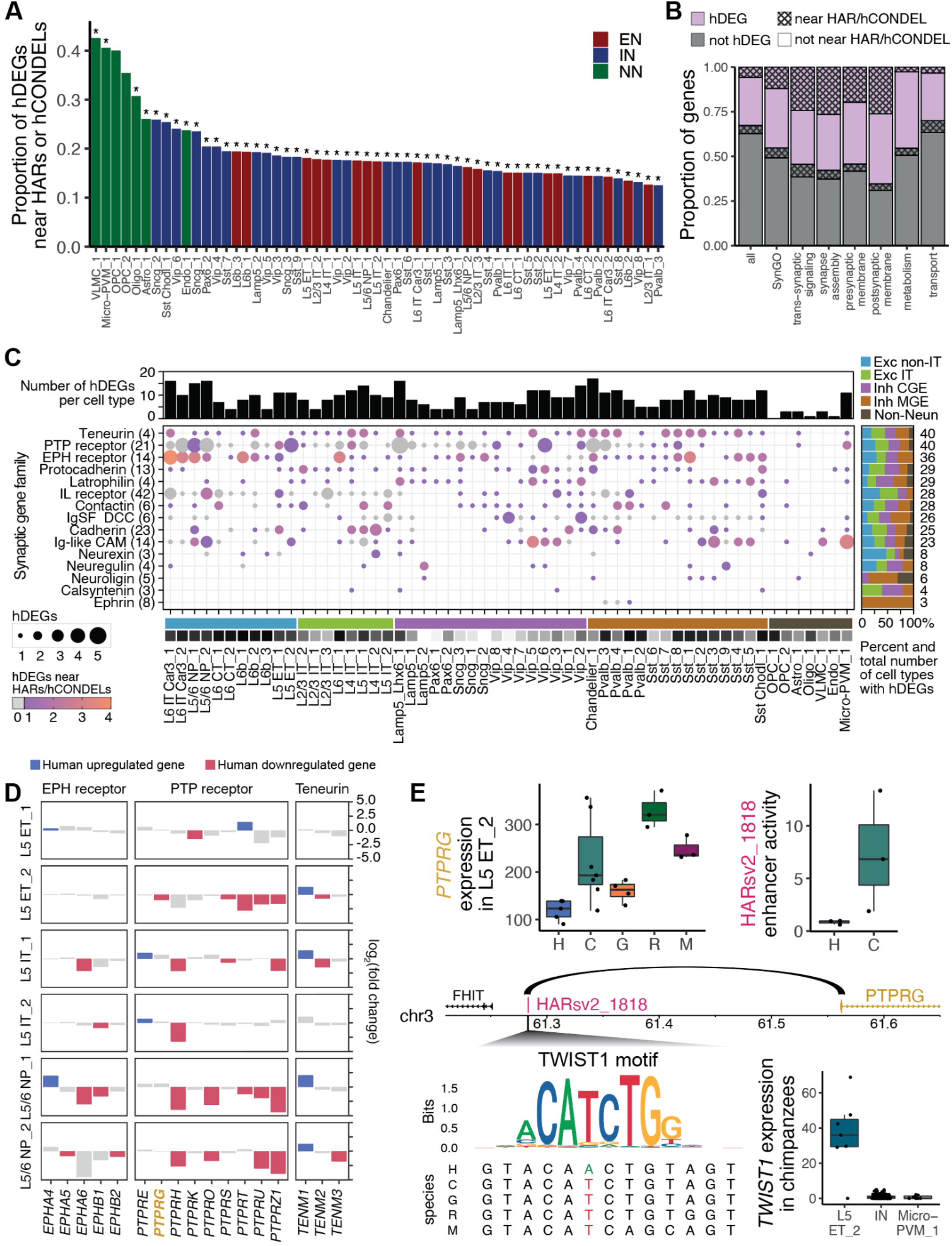
HARs and hCONDELs are enriched near hDEGs. **(A)** HARs and hCONDELs are enriched near hDEGs in virtually all consensus types. Asterisks indicate significance at 5% FDR (Materials and Methods). EN: excitatory neuron, IN: inhibitory neuron, NN: non-neuronal. **(B)** hDEGs, as well as hDEGs near HARs or hCONDELs, are enriched for genes annotated in SynGO (P < 10^−10^) (*25*), particularly those involved in trans-synaptic signaling (P < 0.001) and synapse assembly (P < 0.006), or located at the presynaptic membrane (P < 0.003) and postsynaptic membrane (P < 0.01). **(C)** Synaptic cell-adhesion families with highly divergent expression patterns. Dot sizes represent the number of hDEGs per family in consensus types, and the color represent the number of those hDEGs that are near HARs or hCONDELs (gray indicates that hDEGs are not near any HARs or hCONDELs). Squares above the consensus type labels indicate the average laminar depth of that type from layer 1 (white) to layer 6 (black). **(D)** Patterns of expression change between humans and NHPs for three highly-divergent families in consensus types of excitatory neurons located in L5. **(E)** *PTPRG* has decreased expression in human L5 ET_2 (each point represents normalized pseudobulk gene expression per individual). Its promoter interacts with the intergenic region containing HARsv2_1818 (*15*), which has decreased enhancer activity in human SH-SY5Y cells (*13*). A base pair change in the human HARsv2_1818 sequence removes a potential binding site for *TWIST1*, which is highly expressed in L5 ET_2.

Since hDEGs are highly enriched for synaptic genes (**Fig. 5H**), we asked whether a subset of hDEGs that are potentially adaptive (i.e. near HARs or hCONDELs) are associated with specialized localizations or molecular functions of the synapse by performing gene set enrichment analysis using SynGO (*25*). We found a significant enrichment of hDEGs among SynGO genes compared to all expressed genes (P < 10^−16^) and a further enrichment of hDEGs near HARs and hCONDELs within SynGO genes (P < 10^−10^) (**Fig. 6B; Table S7**). SynGO terms related to synapse assembly, synaptic membrane organization, and *trans*-synaptic signaling were the most enriched, while other terms, including synaptic transport, metabolism, cytoskeleton, and vesicle exocytosis machinery, were not significantly enriched (**Fig. 6B; Fig. S16; Fig. S17; Table S7**). These results suggest that divergent expression patterns in humans compared to NHPs are mainly associated with genetic programs involved in the formation of synaptic connections. Consistent with this idea, we observed an enrichment for hDEGs within gene families that establish ligand-receptor interactions involved in cellular and synaptic adhesion functions (P < 10^−20^) (**Fig. 6C; Fig. S18A,B**). Notably, hDEGs near HARs and hCONDELs are also enriched among genes encoding synaptic adhesion molecules (P < 10^−6^) (**Fig. 6C; Fig. S18; Table S7**). We also found that hDEGs, including those near HARs and hCONDELs, are enriched in other gene families encoding for cell-surface ligand-receptor pairs involved in growth signaling (**Fig. S18; Table S7**). In contrast, hDEGs near HARs and hCONDELs are not enriched in gene families encoding cytoskeletal, scaffolding, and intracellular regulatory components (**Fig. S18; Fig. S19**). Altogether, these findings suggest that genes that show divergent expression in human cortical cells are predominantly related to molecular processes involved in the assembly and specification of synapses.

We next asked how synaptic genetic programs have been differentially used across evolution in specific consensus types. Strikingly, three families of synaptic adhesion molecules (teneurins, PTP and EPH receptors) include hDEGs in almost all consensus types (**Fig. 6C**), while most gene families changed only in a subset of types (**Fig. S18; Fig. S19**). The median number of hDEGs within synaptic adhesion gene families is 20 genes, with a few consensus types (10 out of 57) containing more than 30 hDEGs in these gene families (**Fig. 6C; Fig. S18B**). These synaptically divergent cell types include interneurons with highly specialized connectivity, such as Chandelier and Lamp5_Lhx6 consensus types and several types of deep-layer excitatory neurons (**Fig. S18B**). Moreover, some gene families (neurexins, interleukin receptors, FLRT proteins, and Trk receptors) mainly changed in excitatory types in deep cortical layers, while other families (neuroligins, protocadherins, latrophilins, and Ig superfamily DCC receptors) primarily changed in inhibitory types (**Fig. 6C; Fig. S18**). Interestingly, Pvalb interneurons and deep layer excitatory neurons are known to establish specific microcircuits in deeper cortical layers (*46*), and those types show complementary expression changes in ephrin ligands and receptors, respectively (**Fig. 6C; Fig. S18B**). This suggests possible cell type-specific transcriptional reorganization during human cortical evolution that may underlie differences in synaptic connectivity in specific microcircuits.

Analysis of specific members within synaptic adhesion families revealed a surprising level of differential usage of hDEGs within cellular neighborhoods. There are multiple hDEGs within the PTP receptors, EPH receptors, and teneurins among L5-6 excitatory neurons (**Fig. 6D; Fig. S20A,B**). Twelve genes within these families (*EPHA3, EPHA4, EPHA5, EPHA7, EPHB6, PTPRF, PTPRG, PTPRHK, PTPRS, PTPRT, PTPRU, TENM3*) changed their expression in only 1-2 consensus types within the 14 consensus types of L5-6 excitatory neurons (**Fig. S18B**). Similarly, several genes that only diverge in expression in inhibitory interneurons also showed selective changes in only 1-2 consensus types (*CDH1, CDH2, CDH24, EFNA5, EFNB2, IGSF9B, LGI1, LGI2, LRFN5, SLITRK4*) (**Fig. S18B; Fig. S20C**). Taken together, our data highlight human specializations of synaptic gene programs that are highly localized to specific cell types.

We leveraged existing data to identify human-specific sequence changes in regulatory regions linked to hDEGs that may drive differential expression in select cell types. For example, *PTPRG* is a member of the PTP receptor family that acts as presynaptic organizers for synapse assembly (*47*). Genetic variants in *PTPRG* have been associated with neuropsychiatric disorders, and *Ptprg* mutant mice show memory deficits, supporting an important role for *PTPRG* in cognitive function (*48–51*). We found that *PTPRG* was widely expressed across cell types (**Fig. S21A**) and had lower expression in human compared to NHPs in four consensus types: one excitatory neuron type (L5 ET_2), microglia (Micro-PVM_1), and two inhibitory neuron types (Vip_2 and Vip_6) (**Fig. 6E; Fig. S21B**). *PTPRG* is located near HARv2_1818 (chr3:61283266-61283416, hg38), which has decreased enhancer activity from the human sequence compared to the orthologous chimpanzee sequence in reporter assays in a human neuronal cell line (*13*). Intriguingly, a 5 kb genomic interval that includes HARsv2_1818 has been shown to interact with the promoter of *PTPRG*, specifically in excitatory neurons, but not in inhibitory neurons or microglia (*14, 15*). This raises the possibility that decreased enhancer activity from HARsv2_1818 in humans may have decreased *PTPRG* expression specifically in the excitatory neuron consensus type L5 ET_2 and that separate regulatory mechanisms may decrease *PTPRG* expression in microglia and specific inhibitory neuron consensus types. In support of this hypothesis, there is a base pair substitution in the human HARsv2_1818 sequence that removes a binding site for TWIST1, a basic helix-loop-helix transcription factor. We find that *TWIST1* is expressed predominantly in L5 ET_2 compared to microglia or inhibitory neuron consensus types (**Fig. 6E; Fig. S21C**), further suggesting that human-specific sequence changes in HARsv2_1818 may specifically decrease *PTPRG* expression in L5 ET_2. We extended this analysis to link 112 HARs to 92 hDEGs in neurons using existing data (*14, 15*), and we posit that genomic interaction data from specific cell types may reveal additional genes that may be regulated by human-specific sequence changes.

## Discussion

Transcriptomic profiling of over 570,000 nuclei from the MTG region of primate neocortex reveals a remarkably conserved cellular architecture across humans and four NHPs: chimpanzees, gorillas, macaques, and marmosets. Humans and the other great apes have nearly identical proportions and laminar distributions of consensus types, while marmosets are the most distinct with markedly increased proportions of L5 ET and L6 IT Car3 excitatory neurons and chandelier interneurons. Great apes have equal proportions of two major subtypes of L6 IT Car3 neurons that have high or low *CUX2* expression and distinct positions in layers 5 and 6. Curiously, macaques had mostly Low-CUX2 neurons, and marmosets had mostly High-*CUX2* neurons; this leaves the ancestral state among primates unresolved. In contrast, L6 IT Car3 neurons in mice express markers of both subtypes and are transcriptomically homogeneous across the cortex although they project to diverse cortical targets including proximal areas and homotypic areas in the contralateral hemisphere (*52*). Remarkably, High-*CUX2* neurons are selectively enriched in language-related regions in the human temporal and parietal cortex (MTG, primary auditory cortex, and angular gyrus), and these neurons may have distinct connectivity and contribute to the functional specializations of these regions.

Gene expression differences are more pronounced than cell type proportion differences across primates, including great apes. These differences mostly parallel evolutionary distances. One exception is that neuronal expression in chimpanzees resembles gorillas more than humans, and this is consistent with faster divergence along the human lineage (*53*). Relative to neurons, evolutionary expression changes are accelerated in microglia, astrocytes, and oligodendrocytes in all primates. In addition, human glia have more highly divergent genes than chimpanzee or gorilla glia, suggesting faster divergence of human microglia and astrocytes as well as oligodendrocytes (*8*) among great apes. Finally, human-specific changes in transcription also involve substantial switching of isoform usage in genes that, surprisingly, often have conserved expression levels.

Humans and NHPs have hundreds of DEGs that are specific to one or a few consensus types and are enriched in molecular pathways related to ribosomal processing, cell connectivity, and synaptic function. Human-specific changes in synaptic gene expression are complex, and distinct families of genes are differentially expressed in select neuronal and non-neuronal types. For example, ephrin molecules specifically differ in PVALB inhibitory cell types, while their cognate receptors (EPH receptors) are changing prominently in deep layer excitatory neurons. Importantly, ephrin/EPH receptor signaling has been shown to promote synaptogenesis in the mouse developing cortex (*54, 55*). Since PVALB interneurons and excitatory neurons form selective patterns of connectivity in a cell type-specific fashion (*46*), the differential expression of ephrins and EPH receptors in these cells observed in primate species could reflect species differences in the formation of inhibitory microcircuits involving specific subtypes of PVALB interneurons and excitatory neurons.

Of note, a substantial proportion of synaptic cell-adhesion genes showed down-regulated expression in human neurons, particularly in gene families encoding PTP receptors, including PTPRG, and EPH receptors (**Fig. 6D**). Some studies have proposed roles in synapse elimination for members of highly-divergent synaptic families, including Pcdh10, ephrin-B1, and ephrin-A2 (*56, 57*). In such a case, reduced expression of negative regulators of synaptic assembly in human neurons could lead to an enhanced ability to form synaptic connections, potentially underlying the greater number of synapses per neuron observed in human cortex compared to NHPs (*58, 59*).

Emerging evidence demonstrates the critical role that non-neuronal cell types play in cortical development, network function, and behavior (*60–65*). Previous molecular assays have identified a role for ErbB4-mediated signaling in astrogenesis, astrocyte-neuron communication, and astrocyte-induced neuronal remodeling, potentially through both paracrine and autocrine signaling (*66–68*). Interestingly, we observed changes in expression of *ERBB4* receptor and its cognate ligands *NRG2* and *NRG3* in human astrocytes compared to chimpanzees and gorillas. Altogether, these findings point towards finely regulated molecular specializations underlying neuronal and glial communication in the human cortex. Our data also serves as a resource for future investigation of human-enriched astrocyte and microglia gene programs associated with disease.

Cell type-specific evolutionary changes in gene expression are likely driven by sequence changes to regulatory regions that can be active with high spatial and temporal precision. This is supported by prior studies of genome sequence evolution in humans and other species that estimated that more than 80% of adaptive sequence change is likely regulatory (*44, 69, 70*). Indeed, we find that previously identified genomic regions that have human-specific sequence changes, such as HARs and hCONDELs, are enriched near hDEGs. Intriguingly, this association is observed for both neuronal and non-neuronal consensus types. In addition to well-described changes in the number and function of neurons in the human brain (*71*), many non-neuronal cell types also undergo comparable changes in the human lineage (*72, 73*).

Moreover, hDEGs, including those near HARs and hCONDELs, have been found to play critical roles in synapse establishment, elimination, and maintenance when expressed by neuronal and non-neuronal cells (*74*). Associating HARs and hCONDELs with hDEGs provides a framework to link regulatory sequence changes to human-specific cellular and circuit-level phenotypes via expression changes in select cell types.

## Methods

### Tissue specimens from primate species

#### Human postmortem tissue specimens

De-identified postmortem adult human brain tissue was obtained after receiving permission from the deceased’s next-of-kin. Tissue collection was performed in accordance with the provisions of the United States Uniform Anatomical Gift Act of 2006 described in the California Health and Safety Code section 7150 (effective 1/1/2008) and other applicable state and federal laws and regulations. The Western Institutional Review Board reviewed tissue collection procedures and determined that they did not constitute human subjects research requiring institutional review board (IRB) review.

Male and female individuals 18–68 years of age with no known history of neuropsychiatric or neurological conditions were considered for inclusion in the study. Routine serological screening for infectious disease (HIV, Hepatitis B, and Hepatitis C) was conducted using individual blood samples and individuals testing positive for infectious disease were excluded from the study. Specimens were screened for RNA quality and samples with average RNA integrity (RIN) values ≥7.0 were considered for inclusion in the study. Postmortem brain specimens were processed as previously described (*16*) (<u>dx.doi.org/10.17504/protocols.io.bf4ajqse</u>). Briefly, coronal brain slabs were cut at 1 cm intervals, frozen in dry-ice cooled isopentane, and transferred to vacuum-sealed bags for storage at −80°C until the time of further use. To isolate the MTG, tissue slabs were briefly transferred to −20°C and the region of interest was removed and subdivided into smaller blocks on a custom temperature controlled cold table. Tissue blocks were stored at −80°C in vacuum-sealed bags until later use.

#### Chimpanzee and gorilla tissue specimens

Chimpanzee tissue was obtained from the National Chimpanzee Brain Resource (supported by NIH grant NS092988). Gorilla samples were collected postmortem following naturally occurring death or euthanasia of the animals for medical conditions at various zoos. Gorilla and chimpanzee brains were divided into 2 cm coronal slabs, flash frozen using dry-ice cooled isopentane, liquid nitrogen, or a −80°C freezer, and then stored in freezer bags at −80°C. Tissues from the MTG were removed from appropriate slabs which were maintained on dry-ice during dissection and were shipped to the Allen Institute overnight on dry-ice.

#### Macaque tissue specimens

Macaque tissue samples were obtained from the University of Washington National Primate Resource Center under a protocol approved by the University of Washington Institutional Animal Care and Use Committee. Immediately following euthanasia, macaque brains were removed and transported to the Allen Institute in artificial cerebral spinal fluid equilibrated with 95% O_2_ and 5% CO_2_. Upon arrival at the Allen Institute, brains were divided down the midline and each hemisphere was subdivided coronally into 0.5 cm slabs. Slabs were flash frozen in dry-ice cooled isopentane, transferred to vacuum-sealed bags, and stored at −80°C. MTG was removed from brain slabs as described above for human tissues.

#### Marmoset tissue specimens

Marmoset experiments were approved by and in accordance with Massachusetts Institute of Technology IACUC protocol number 051705020. Adult marmosets (1.5–2.5 years old, 3 individuals) were deeply sedated by intramuscular injection of ketamine (20–40 mg kg−1) or alfaxalone (5–10 mg kg−1), followed by intravenous injection of sodium pentobarbital (10–30 mg kg−1). When the pedal with-drawal reflex was eliminated and/or the respiratory rate was diminished, animals were trans-cardially perfused with ice-cold sucrose-HEPES buffer (*75*). Whole brains were rapidly extracted into fresh buffer on ice. Sixteen 2-mm coronal blocking cuts were rapidly made using a custom-designed marmoset brain matrix. Slabs were transferred to a dish with ice-cold dissection buffer (*75*). All regions were dissected using a marmoset atlas as reference (*76*), and were snap-frozen in liquid nitrogen or dry ice-cooled isopentane, and stored in individual microcentrifuge tubes at −80 °C.

Temporal lobe dissections targeted area TE3 and TPO on the lateral temporal surface. Though a true homology to catarhine MTG may not exist in marmosets, these areas in marmoset form part of the temporal lobe association cortex. Moreover, on the basis of tract tracing connectivity studies (*77*), TE3 and TPO participate in the ‘default mode network,’ a functionally coupled network of higher-order association cortex that includes MTG in other species (*78*). Cortical area DLPFC targeted the dorsolateral surface of PFC, approximately 2.5-3 mm from the frontal pole. ACC/PFCm included medial frontal cortex anterior to the genu of the corpus callosum. M1 dissections were stained with fluorescent Nissl and targeted the hand/trunk region. S1 like sampled all primary somatosensory areas (A3,A1/2). A1 dissections targeted primary auditory area but likely include some rostral and caudal parabelt cortex. V1 dissections were collected on the dorsal bank of the calcarine sulcus approximately 4-6 mm from the posterior pole.

### Tissue processing and single nucleus RNA-sequencing

#### SMART-seq v4 nucleus isolation and sorting

Vibratome sections of MTG blocks were stained with fluorescent Nissl permitting microdissection of individual cortical layers as previously described (<u>dx.doi.org/10.17504/protocols.io.bq6ymzfw</u>). Nucleus isolation was performed as described (<u>dx.doi.org/10.17504/protocols.io.ztqf6mw</u>). Briefly, single nucleus suspensions were stained with DAPI (4’,6-diamidino-2-phenylindole dihydrochloride, ThermoFisher Scientific, D1306) at a concentration of 0.1μg/ml. Controls were incubated with mouse IgG1k-PE Isotype control (BD Biosciences, 555749, 1:250 dilution) or DAPI alone. To discriminate between neuronal and non-neuronal nuclei, samples were stained with mouse anti-NeuN conjugated to PE (FCMAB317PE, EMD Millipore) at a dilution of 1:500.

Single-nucleus sorting was carried out on either a BD FACSAria II SORP or BD FACSAria Fusion instrument (BD Biosciences) using a 130 μm nozzle and BD Diva software v8.0. A standard gating strategy based on DAPI and NeuN staining was applied to all samples as previously described (*16*). Doublet discrimination gates were used to exclude nuclei multiplets. Individual nuclei were sorted into 96-well plates, briefly centrifuged at 1000 rpm, and stored at −80°C.

#### SMART-seq v4 RNA-sequencing

The SMART-Seq v4 Ultra Low Input RNA Kit for Sequencing (Takara #634894) was used per the manufacturer’s instructions. Standard controls were processed with each batch of experimental samples as previously described. After reverse transcription, cDNA was amplified with 21 PCR cycles. The NexteraXT DNA Library Preparation (Illumina FC-131-1096) kit with NexteraXT Index Kit V2 Sets A-D (FC-131-2001, 2002, 2003, or 2004) was used for sequencing library preparation. Libraries were sequenced on an Illumina HiSeq 2500 instrument (Illumina HiSeq 2500 System, RRID:SCR_016383) using Illumina High Output V4 chemistry. The following instrumentation software was used during data generation workflow; SoftMax Pro v6.5; VWorks v11.3.0.1195 and v13.1.0.1366; Hamilton Run Time Control v4.4.0.7740; Fragment Analyzer v1.2.0.11; Mantis Control Software v3.9.7.19.

#### SMART-seq v4 gene expression quantification

For human, raw read (fastq) files were aligned to the GRCh38 genome sequence (Genome Reference Consortium, 2011) with the RefSeq transcriptome version GRCh38.p2 (RefSeq, RRID:SCR_003496, current as of 4/13/2015) and updated by removing duplicate Entrez gene entries from the gtf reference file for STAR processing, as previously described (*16*). For chimpanzee and gorilla, the Clint_PTRv2 and Susie3 NCBI reference genomes were used for alignment, respectively. For alignment, Illumina sequencing adapters were clipped from the reads using the fastqMCF program (from ea-utils). After clipping, the paired-end reads were mapped using Spliced Transcripts Alignment to a Reference (STAR v2.7.3a, RRID:SCR_015899) using default settings. Reads that did not map to the genome were then aligned to synthetic construct (i.e. ERCC) sequences and the *coli* genome (version ASM584v2). Quantification was performed using summerizeOverlaps from the R package GenomicAlignments v1.18.0. Expression levels were calculated as counts per million (CPM) of exonic plus intronic reads.

#### 10x Chromium RNA-sequencing and expression quantification

Nucleus isolation for 10x Chromium snRNAseq was conducted as described (<u>dx.doi.org/10.17504/protocols.io.y6rfzd6</u>). Gating was as described for SSv4 above. NeuN+ and NeuN-nuclei were sorted into separate tubes and were pooled at a defined ratio (90% NeuN+, 10% NeuN-) after sorting. Sorted samples were centrifuged, frozen in a solution of 1X PBS, 1% BSA, 10% DMSO, and 0.5% RNAsin Plus RNase inhibitor (Promega, N2611), and stored at −80°C until the time of 10x chip loading. Immediately before loading on the 10x Chromium instrument, frozen nuclei were thawed at 37°C, washed, and quantified for loading as described (<u>dx.doi.org/10.17504/protocols.io.nx3dfqn</u>). Samples were processed using the 10x Chromium Single-Cell 3’ Reagent Kit v3 following the manufacturer’s protocol. Gene expression was quantified using the default 10x Cell Ranger v3 (Cell Ranger, RRID:SCR_017344) pipeline. Reference genomes included the modified genome annotation described above for SMART-seq v4 quantification (human), Clint_PTRv2 (chimpanzee), Susie3 (gorilla), and Mmul_10 (rhesus macaque). Introns were annotated as “mRNA”, and intronic reads were included in expression quantification.

#### Marmoset 10x RNA-seq

Unsorted single-nucleus suspensions from frozen marmoset samples were generated as in (*10*). GEM generation and library preparation followed the manufacturer’s protocol (10X Chromium single-cell 3′ v.3, protocol version #CG000183_ ChromiumSingleCell3′_v3_UG_Rev-A). Raw sequencing reads were aligned to the CJ1700 reference. Reads that mapped to exons or introns were assigned to annotated genes.

### RNA-sequencing processing and clustering

#### Cell type label transfer

Human M1 reference taxonomy subclass labels (*12*) were transferred to nuclei in the current MTG dataset using Seurat’s label transfer (3000 high variance genes using the ‘vst’ method then filtered through exclusion list). For human label mapping to other species, higher variance genes were included from a list of orthologous genes (14,870 genes; downloaded from NCBI Homologene (https://www.ncbi.nlm.nih.gov/homologene) in November 2019; RRID SCR_002924). This was carried out for each species and RNA-seq modality dataset; for example, human-Cv3 and human-SSv4 were labeled independently. Each dataset was subdivided into 5 neighborhoods – IT and Non-IT excitatory neurons, CGE- and MGE-derived interneurons, and non-neuronal cells – based on marker genes and transferred subclass labels from published studies of human and mouse cortical cell types and cluster grouping relationships in a reduced dimensional gene expression space.

#### Filtering low-quality nuclei

SSv4 nuclei were included for analysis if they passed all QC criteria:

> 30% cDNA longer than 400 base pairs
> 500,000 reads aligned to exonic or intronic sequence
> 40% of total reads aligned
> 50% unique reads
> 0.7 TA nucleotide ratio

QC was then performed at the neighborhood level. Neighborhoods were integrated together across all species and modality; for example, deep excitatory neurons from human-Cv3, human-SSv4, Chimp-Cv3, etc. datasets were integrated using Seurat integration functions with 2000 high variance genes from the orthologous gene list. Integrated neighborhoods were Louvain clustered into over 100 meta cells, and Low-quality meta cells were removed from the dataset based on relatively low UMI or gene counts (included glia and neurons with greater than 500 and 1000 genes detected, respectively), predicted doublets (include nuclei with doublet scores under 0.3), and/or subclass label prediction metrics within the neighborhood (ie excitatory labeled nuclei that clustered with majority inhibitory or non-neuronal nuclei).

#### RNA-seq clustering

Nuclei were normalized using SCTransform (*18*), and neighborhoods were integrated together within a species and across individuals and modalities by identifying mutual nearest neighbor anchors and applying canonical correlation analysis as implemented in Seurat (*17*). For example, deep excitatory neurons from human-Cv3 were split by individual and integrated with the human-SSv4 deep excitatory neurons. Integrated neighborhoods were Louvain clustered into over 100 meta cells. Meta cells were then merged with their nearest neighboring meta cell until merging criteria were sufficed, a split and merge approach that has been previously described (*12*). The remaining clusters underwent further QC to exclude Low-quality and outlier populations. These exclusion criteria were based on irregular groupings of metadata features that resided within a cluster.

### Robustness tests of cell subclasses using MetaNeighbor

MetaNeighbor v1.12 (*38, 39*) was used to provide a measure of neuronal and non-neuronal subclass and cluster replicability within and across species. We subset snRNA-seq datasets from each species to the list of common orthologs before further analysis. For each assessment, we identified highly variable genes using the get_variable_genes function from MetaNeighbor. In order to identify homologous cell types, we used the MetaNeighborUS function, with the fast_version and one_vs_best parameters set to TRUE. The one_vs_best parameter identifies highly specific cross-dataset matches by reporting the performance of the closest neighboring cell type over the second closest as a match for the training cell type, and the results are reported as the relative classification specificity (AUROC). This step identified highly replicable cell types within each species and across each species pair. All 24 subclasses are highly replicable within and across species (one_vs_best AUROC of 0.96 within species and 0.93 across species in Fig. S4A).

### Defining cross-species consensus cell types

While cell type clusters are highly replicable within each species (one_vs_best AUROC of 0.93 for neurons and 0.87 for non-neurons), multiple transcriptionally similar clusters mapped to each other across each species pair (average cross-species one_vs_best AUROC of 0.76). To build a consensus cell type taxonomy across species, we defined a cross-species cluster as a group of clusters that are either reciprocal best hits or clusters with AUROC > 0.6 in the one_vs_best mode in at least one pair of species. This lower threshold (AUROC>0.6) reflects the high difficulty/specificity of testing only against the best performing other cell type. We identified 86 cross-species clusters, each containing clusters from at least two primates. Any unmapped clusters were assigned to one of the 86 cross-species clusters based on their transcriptional similarity. For each unmapped cluster, the top 10 of their closest neighbors are identified using MetaNeighborUS one_vs_all cluster replicability scores, and the unmapped cluster is assigned to the cross-species cluster in which a strict majority of its nearest neighbors belong. For clusters with no hits, this is repeated using the top 20 closest neighbors, still requiring a strict majority to assign a cross-species type. 594 clusters present in five primates (i.e., union) mapped to 86 cross-species clusters, with 493 clusters present across 57 consensus cross-species clusters shared by all five primates.

### Cell type taxonomy generation

For each species, a taxonomy was built using the final set of clusters and was annotated using subclass mapping scores, dendrogram relationships, marker gene expression, and inferred laminar distributions. Within-species taxonomy dendrograms were generated using build_dend function from scrattch_hicat R package. A matrix of cluster median log2(cpm + 1) expression across the 3000 High-variance genes for Cv3 nuclei from a given species were used as input. The cross-species dendrogram was generated with a similar workflow but was downsampled to a maximum of 100 nuclei per cross-species cluster per species. The 3000 High-variance genes used for dendrogram construction were identified from the downsampled matrix containing Cv3 nuclei from all five species. We generated the complete cross-species cluster dendrogram using average-linkage hierarchical clustering with (1 - average MetaNeighborUS one_vs_all cluster replicability scores) for each pair of 86 cross-species clusters as a measure of distance between cell types. The consensus cross-species taxonomy is visualized (in **Fig. 5E**) by pruning the complete dendrogram to retain only the 57 homologous cell types shared by all five species.

### Cell type comparisons across species

#### Differential gene expression

To identify subclass marker genes within a species, Cv3 datasets from each species were downsampled to a maximum of 100 nuclei per cluster per individual. Differentially expressed marker genes were then identified using the FindAllMarkers function from Seurat, using the Wilcoxon sum rank test on log-normalized matrices with a maximum of 500 nuclei per group (subclass vs. all other nuclei as background). Statistical thresholds for markers are indicated in their respective figures. To identify species marker genes across subclasses, Cv3 datasets from each species were downsampled to a maximum of 50 nuclei per cluster per individual. Downsampled counts matrices were then grouped into pseudo-bulk replicates (species, individual, subclass) and the counts were summed per replicate. DESeq2 functionality was then used to perform a differential expression analysis between species pairs (or comparisons of interest) for each subclass using the Wald test statistic.

#### Expression correlations

Subclasses were compared between each pair of species using Spearman correlations on subclass median log2(cpm + 1) expression of orthologous genes that had a median value greater than 0 in both species. These Spearman correlations were then visualized as heatmaps and also compared to the human-centric evolutionary distance from each species in Figure 2. Similarly, subclasses were compared across individuals within each species, and the average Spearman correlation of all pairwise comparisons of individuals was calculated.

#### Taxonomy comparisons

To assess homologies between clusters from taxonomies of different species or different studies, we constructed Euclidean distance heatmaps that were anchored on one side by the taxonomies’ dendrogram. The heatmaps display the cluster labels of a single taxonomy on either end, and the heatmap values represent the Euclidean distance between cluster centroids in the reduced dimensional space using 30-50 principal components from a PC analysis. In the case of cross-species comparisons, the reduced space was derived from Cv3 data. The −log(Euclidean distance) is plotted, with smaller values indicating more similar transcriptomic profiles.

#### Estimating differential isoform usage between great apes

We used Smart-seq single-nucleus RNA-seq data from humans (∼14,500 cells), chimpanzees (∼3,500 cells), and gorillas (∼4,300 cells) to assess isoform switching between the species for each cell subclass. The RNA-seq reads were mapped to each species’ genome using STAR as described above. The isoforms were quantified using RSEM on a common set of annotated transcripts (TransMap V5 downloaded from the UCSC browser, RRID:SCR_005780) by aggregating reads from cells in each cell subtype using a pseudo-bulk method:

1. Aggregated reads from cells in each subclass
2. Mapped reads to the human, chimpanzee, or gorilla reference genome with STAR 2.7.7a using default parameters
3. Transformed genomic coordinates into transcriptomic coordinates using STAR parameter: -- quantMode TranscriptomeSAM
4. Quantified isoform and gene expression using RSEM v1.3.3 parameters (RSEM, RRID:SCR_013027): --bam --seed 12345 --paired-end --forward-prob 0.5 --single-cell-prior --calc-ci

The isoform proportion metric (isoP) was defined as the isoform expression (transcripts per million, TPM) normalized by the total expression of the gene the isoform belongs to. To focus on highly expressed genes, we considered only isoforms originating from the top 50% (ranked by gene expression) of genes for each species. To control the variability of isoP values, we derived the 80% confidence intervals by comparing the isoP-values of different donors for each species using the following procedure:

1. The isoP values (ranging from 0 to 1) for donor 1 are binned into 10 bins of size 0.1.
2. The isoforms in each bin are sorted by the isoP values in donor 2.
3. The lower and upper bounds of the 80% isoP confidence interval are defined as 10% and 90% percentile of this sorted list.
4. The procedure was repeated, switching donors 1 and 2, and the isoP confidence interval bounds values from the two calculations were averaged.

The isoform switching between species was considered significant for isoforms whose confidence intervals were non-overlapping. We defined cross-species isoform switches as those that involved a major isoform in one of the species (i.e., isoP > 0.5) and report them in **Table S4**. A subset of isoforms with strong cross-species switching were identified that had isoP > 0.7 in one species, isoP < 0.1 in the other species, and >3-fold change in proportions between the species.

#### Identifying changes in cell type proportions across species

Cell type proportions are compositional, where the gain or loss of one population necessarily affects the proportions of the others, so we used scCODA (*42*) to determine which changes in cell class, subclass, and cluster proportions across species were statistically significant. We focused these analyses on neuronal populations since these were deeply sampled in all five species based on sorting of nuclei with NeuN immunostaining. The proportion of each neuronal class, subclass, and cluster was estimated using a Bayesian approach where proportion differences across individuals were used to estimate the posterior. All compositional and categorical analyses require a reference population to describe differences with respect to and, because we were uncertain which populations should be unchanged, we iteratively used each cell type and each species as a reference when computing abundance changes. To account for sex differences, we included it as a covariate when testing for abundance changes. We report the effect size of each species and sex for each cell subclass and used a mean inclusion probability cutoff of 0.7 for calling a population credibly different.

### *In situ* profiling of gene expression

#### MERFISH data collection

Human postmortem frozen brain tissue was embedded in Optimum Cutting Temperature medium (VWR,25608-930) and sectioned on a Leica cryostat at −17 C at 10 um onto Vizgen MERSCOPE coverslips (VIZGEN 2040003). These sections were then processed for MERSCOPE imaging according to the manufacturer’s instructions. Briefly: sections were allowed to adhere to these coverslips at room temperature for 10 min prior to a 1 min wash in nuclease-free phosphate buffered saline (PBS) and fixation for 15 min in 4% paraformaldehyde in PBS. Fixation was followed by 3×5 minute washes in PBS prior to a 1 min wash in 70% ethanol. Fixed sections were then stored in 70% ethanol at 4 C prior to use and for up to one month. Human sections were photobleached using a 150W LED array for 72 h at 4 C prior to hybridization then washed in 5 ml Sample Prep Wash Buffer (VIZGEN 20300001) in a 5 cm petri dish. Sections were then incubated in 5 ml Formamide Wash Buffer (VIZGEN 20300002) at 37 C for 30 min. Sections were hybridized by placing 50 ul of VIZGEN-supplied Gene Panel Mix onto the section, covering with parafilm and incubating at 37 C for 36-48 h in a humidified hybridization oven.

Following hybridization, sections were washed twice in 5 ml Formamide Wash Buffer for 30 min at 47 C. Sections were then embedded in acrylamide by polymerizing VIZGEN Embedding Premix (VIZGEN 20300004) according to the manufacturer’s instructions. Sections were embedded by inverting sections onto 110 ul of Embedding Premix and 10% Ammonium Persulfate (Sigma A3678) and TEMED (BioRad 161-0800) solution applied to a Gel Slick (Lonza 50640) treated 2×3 glass slide. The coverslips were pressed gently onto the acrylamide solution and allowed to polymerize for 1.5 h. Following embedding, sections were cleared for 24-48 h with a mixture of VIZGEN Clearing Solution (VIZGEN 20300003) and Proteinase K (New England Biolabs P8107S) according to the Manufacturer’s instructions. Following clearing, sections were washed twice for 5 min in Sample Prep Wash Buffer (PN 20300001). VIZGEN DAPI and PolyT Stain (PN 20300021) was applied to each section for 15 min followed by a 10 min wash in Formamide Wash Buffer. Formamide Wash Buffer was removed and replaced with Sample Prep Wash Buffer during MERSCOPE set up. 100 ul of RNAse Inhibitor (New England BioLabs M0314L) was added to 250 ul of Imaging Buffer Activator (PN 203000015) and this mixture was added via the cartridge activation port to a pre-thawed and mixed MERSCOPE Imaging cartridge (VIZGEN PN1040004). 15 ml mineral oil (Millipore-Sigma m5904-6×500ML) was added to the activation port and the MERSCOPE fluidics system was primed according to VIZGEN instructions. The flow chamber was assembled with the hybridized and cleared section coverslip according to VIZGEN specifications and the imaging session was initiated after collection of a 10X mosaic DAPI image and selection of the imaging area. For specimens that passed minimum count threshold, imaging was initiated and processing completed according to VIZGEN proprietary protocol.

The 140 gene Human cortical panel was selected using a combination of manual and algorithmic based strategies requiring a reference single cell/nucleus RNA-seq data set from the same tissue, in this case the human MTG snRNAseq dataset and resulting taxonomy(Hodge and Bakken 2019). First, an initial set of High-confidence marker genes are selected through a combination of literature search and analysis of the reference data. These genes are used as input for a greedy algorithm (detailed below). Second, the reference RNA-seq data set is filtered to only include genes compatible with mFISH. Retained genes need to be 1) long enough to allow probe design (> 960 base pairs); 2) expressed highly enough to be detected (FPKM >= 10), but not so high as to overcrowd the signal of other genes in a cell (FPKM < 500); 3) expressed with low expression in off-target cells (FPKM < 50 in non-neuronal cells); and 4) differentially expressed between cell types (top 500 remaining genes by marker score20). To more evenly sample each cell type, the reference data set is also filtered to include a maximum of 50 cells per cluster.

Cell type mapping in MERSCOPE data: Any genes not matched across both the MERSCOPE gene panel and the snRNASeq mapping taxonomy were filtered from the snRNASeq dataset. We calculated the mean gene expression for each gene in each snRNAseq cluster. We assigned MERSCOPE cells to snRNAseq clusters by finding the nearest cluster to the mean expression vectors of the snRNASeq clusters using the cosine distance. All scripts and data used are available at: https://github.com/AllenInstitute/Great_Ape_MTG.

The main step of gene selection uses a greedy algorithm to iteratively add genes to the initial set. To do this, each cell in the filtered reference data set is mapped to a cell type by taking the Pearson correlation of its expression levels with each cluster median using the initial gene set of size n, and the cluster corresponding to the maximum value is defined as the “mapped cluster”. The “mapping distance” is then defined as the average cluster distance between the mapped cluster and the originally assigned cluster for each cell. In this case a weighted cluster distance, defined as one minus the Pearson correlation between cluster medians calculated across all filtered genes, is used to penalize cases where cells are mapped to very different types, but an unweighted distance, defined as the fraction of cells that do not map to their assigned cluster, could also be used. This mapping step is repeated for every possible n+1 gene set in the filtered reference data set, and the set with minimum cluster distance is retained as the new gene set. These steps are repeated using the new get set (of size n+1) until a gene panel of the desired size is attained. Code for reproducing this gene selection strategy is available as part of the mfishtools R library (https://github.com/AllenInstitute/mfishtools).

#### RNAscope

Fresh-frozen human postmortem brain tissues were sectioned at 16-25 μm onto Superfrost Plus glass slides (Fisher Scientific). Sections were dried for 20 minutes on dry ice and then vacuum sealed and stored at −80°C until use. The RNAscope multiplex fluorescent V2 kit was used per the manufacturer’s instructions for fresh-frozen tissue sections (ACD Bio), except that slides were fixed 60 minutes in 4% paraformaldehyde in 1X PBS at 4°C and treated with protease for 15 minutes. Sections were imaged using a 40X oil immersion lens on a Nikon TiE fluorescence microscope equipped with NIS-Elements Advanced Research imaging software (v4.20, RRID:SCR_014329). Positive cells were called by manual assessment of RNA spots for each gene. Cells were called positive for a gene if they contained ≥ 5 RNA spots for that gene. High versus low expression of *CUX2* was determined by measuring fluorescence intensity for that gene in ImageJ. Lipofuscin autofluorescence was distinguished from RNA spot signal based on the broad fluorescence spectrum and larger size of lipofuscin granules. Staining for each probe combination was repeated with similar results on at least 2 separate individuals and on at least 2 sections per individual. Images were assessed with the FIJI distribution of ImageJ v1.52p and with NIS-Elements v4.20. RNAscope probes used were *CUX2* (ACD Bio, 425581-C3), *LDB2* (1003951-C2), and *SMYD1* (493951-C2).

#### Analysis of great ape species pairwise comparison for glial cells

Significant differential gene expression in pairwise comparisons of glial cells (astrocytes, microglia, oligodendrocytes) across great ape species was determined at log2 fold-change > 0.5 and FDR < 0.01. Among DEGs from great ape pairwise comparisons, species-specific highly divergent genes were identified as having >10 fold change in expression in a given species relative to the other two great ape species, and with a threshold of gene expression of normalized gene counts >5 in at least one species. GO enrichment analysis was performed using the Bioconductor package ‘clusterProfiler’ (https://bioconductor.org/packages/release/bioc/html/clusterProfiler.html), and the Fisher’s exact test was used for SynGO enrichment analysis (https://www.syngoportal.org/). GO and SynGO analyses were performed on the union of DEGs from the pairwise comparison between human and chimpanzee and the pairwise comparison between human and gorilla to increase power to detect significant GO terms. GO terms under biological process, molecular function, and cellular component categories were considered in the analysis. Significance for enriched terms was determined at 5% FDR. All MTG expressed genes in the consensus cell types (astrocytes, microglia, oligodendrocytes) were considered as the background gene set in the respective analyses. Gene expression change in glial cell types shown in heatmaps (Figures 3E; S5A,B,G; S6E,F) is calculated as the log2 ratio of normalized expression counts in a given species relative to the other two great ape species. To analyze astrocyte genes associated with perisynaptic astrocytic processes, a list of genes encoding proteins enriched at astrocyte-neuron junctions was used from a proteomic study in the mouse cortex (*26*). To analyze microglia genes associated with intercellular communication and signaling, a list of genes predicted to act as the ligand-receptor interactome of microglia-neuron communication was used from a recent study in the mouse cortex (*31*).

#### Enrichment of HARs and hCONDELs near hDEGs

The set of HARs used in our analysis was obtained from (*13*). The set of hCONDELs was obtained from (*44, 45*), and only hCONDELs that could be mapped to a syntenic orthologous location in hg38 were retained (1175 total) (*79*). We assigned intronic HARs/hCONDELs to the genes they are intronic to and intergenic HARs/hCONDELs to the closest upstream and downstream genes (**Table S7**) using Ensembl hg38 annotations obtained in May 2021.

63.2% of HARs and 59.4% of hCONDELs are intronic. For 83.2% of the 1165 intergenic HARs, at least one of their assigned genes is within 100kb. For 85.5% of the 477 intergenic hCONDELs, at least one of their assigned genes is within 100kb. To test whether HARs/hCONDELs are enriched near hDEGs in each consensus type, we used the binomial test, which accounts for differences in the size of the regulatory domain for each gene (*80*). The regulatory domain of each gene is defined as the gene body, along with the upstream and downstream intergenic regions that extend to the nearest flanking genes, with an upper bound of 5 Mb in total size. Significance was determined at 5% FDR.

#### SynGO and synaptic gene family enrichment

To analyze the association between synaptic terms and human divergent gene expression patterns, we used an expert-curated database of GO annotations of synapse-related terms known as SynGO (*25*). To test whether hDEGs and hDEGs near HARs/hCONDELs are enriched in SynGO terms, we used Fisher’s exact test. We focused on SynGO terms within the first and second hierarchical levels of SynGO that broadly comprise the entire range of Cellular Components (CC) and Biological Processes (BP) terms, allowing for the visualization of enrichment patterns across a wide range of synaptic localizations and processes (in Figure S16). We grouped SynGO terms into two levels based on their hierarchical organization in SynGO (https://www.syngoportal.org/), corresponding to the following reference codes: 11 terms within level 1 (A1, B1, C1, D1, E1, F1, G1, H1, I1, J1, K1), and 71 terms within level 2 (A2-3, B2-11, C2, D2-11, E2, F2-10, G2-7, H2-4, I2-15, J2-11, K2-6). For synaptic gene families, we examined 15 functionally related categories: (1) families of cell-adhesion and synaptic-adhesion molecules, (2) families of ligand-receptor complexes involved in growth factor signaling, (3) families of other cell-surface receptors and ligands, (4) families of other G protein-coupled receptors (GPCRs) and their ligands (including orphan GPCRs), (5) families of ligand-receptor complexes involved in neuropeptidergic signaling and related GPCRs and ligands, (6) families of neurotransmitter-gated receptors and other ligand-gated receptors (including glutamate ionotropic receptors), (7) Ras GTPase superfamily, (8) families of Ras GAP and GEF signaling molecules, (9) families of other regulatory molecules and structural scaffolding proteins, (10) families related to other signaling complexes including intracellular kinases and phosphatases, (11) families related to the extracellular matrix (ECM) and proteoglycan families, (12) families related to cytoskeletal composition and organization and other related proteins, (13) families involved in synaptic vesicle exocytosis and other membrane fusion components, (14) families of proteases and peptidases, (15) families of voltage-gated ion channels and other gated ion channels and solute transporters. For each of these, we assembled a comprehensive list based on HGNC reference and a previously-curated catalog of synaptic molecules (*81*) (Table S7). Significance was determined at 5% FDR. “All” genes are genes that are expressed and can be assessed for differential expression by DESeq2 in at least one consensus type.

## Acknowledgements

We thank the tissue procurement, tissue processing and facilities teams at the Allen Institute for Brain Science for assistance with the transport and processing of postmortem and neurosurgical brain specimens; the technology team at the Allen Institute for assistance with data management; M. Vawter, J. Davis and the San Diego Medical Examiner’s Office for assistance with postmortem tissue donations. We thank M. Hawrylycz for improvements to taxonomy visualizations. Research reported in this publication was supported by the National Institute Of Mental Health of the National Institutes of Health under Award Numbers U01MH114812 and U19MH114830. The content is solely the responsibility of the authors and does not necessarily represent the official views of the National Institutes of Health. The authors thank the founder of the Allen Institute, Paul G. Allen, for his vision, encouragement and support.

## Funding

Allen Discovery Center for Human Brain Evolution (CAW, DEA, JHS, MB)

Dutch Research Council (NWO) Applied and Engineering Sciences (AES) grant 3DOMICS 17126 (BPL, JE, TH, TK)

Dutch Research Council (NWO) Gravitation Program BRAINSCAPES 024.004.012 (BPL, SB, TH)

EMBO Postdoctoral Fellowship (ALTF 336-2022) (DEA)

Helen Hay Whitney Fellowship (JHS)

NARSAD Young Investigator Award (MC)

National Institutes of Health grant F32MH114501 (MC)

National Institutes of Health grant HG011641 (CCS, WDH)

National Institutes of Health grant NS092988 (CCS, WDH)

National Institutes of Health grant R01 HG009318 (AD)

National Institutes of Health grant R01LM012736 (HS, JGi)

National Institutes of Health grant R01MH113005 (HS, JGi)

National Institutes of Health grant U01MH114812 (AMY, AT, CR, DB, DM, ESL, JGo, KSi, KSm, KW, MT, NLJ, RDH, SD, SL, TEB, TP)

National Institutes of Health grant U01MH114819 (FMK, GF, SAM)

National Institutes of Health grant U19MH114821 (HS, JGi)

National Institutes of Health grant U19MH114830 (AG, AT, DB, DM, JGo, KSm, KW, MT, TP) NSF EF-2021785 (CCS, WDH)

Y. Eva Tan Postdoctoral Fellowship (JHS)

## Author contributions

Sample preparation and RNA data generation: AG, AMY, AT, CCS, CDK, CR, DB, DM, ESL, FMK, GF, JG, KS, KS, KW, MT, RDH, SAM, SD, SL, TEB, TP, WDH

Spatial transcriptomic data generation: BL, DM, EG, JC

Data archive / Infrastructure: JG, SS

Cytosplore Viewer software: BPL, JE, SB, TH, TK

Data analysis: AD, BL, BPL, CAW, DEA, EG, ESL, FMK, HS, JC, JE, JG, JG, JHS, KS, KT, MB, MC, NC, NLJ, SB, TEB, TH

Data interpretation: AD, CAW, CCS, DEA, ESL, FMK, HS, JC, JG, JG, JHS, KT, MB, MC, NC, NLJ, RDH, SD, TEB

Writing manuscript: CAW, CCS, DEA, ESL, JC, JG, JHS, NLJ, RDH, TEB

## Data and materials availability

Raw sequence data were produced as part of the BRAIN Initiative Cell Census Network (BICCN: RRID:SCR_015820) are available for download from the Neuroscience Multi-omics Archive (RRID:SCR_016152; https://assets.nemoarchive.org/dat-net1412/) and the Brain Cell Data Center (RRID:SCR_017266; https://biccn.org/data). Code for analysis and generation of figures is available for download from https://github.com/AllenInstitute/Great_Ape_MTG. Visualization and analysis tools for integrated species comparison are available using Cytosplore Viewer (RRID SCR_018330; https://viewer.cytosplore.org/). These tools allow comparison of gene expression in consensus clusters across species, as well as species-specific clusters and to calculate differential expression within and among species. The following publicly available datasets were used for analysis: Synaptic Gene Ontology (SynGO) and orthologous genes across species from NCBI Homologene (downloaded November 2019). MTG human SMARTseq v4 data (https://portal.brain-map.org/atlases-and-data/rnaseq/human-mtg-smart-seq, https://assets.nemoarchive.org/dat-swzf4kc).

## Supplementary Materials

Figs. 1 to 19

Tables 1 to 7

