## Supplemental Figures 1-21 and Tables 1-7 for "Comparative transcriptomics reveals human-specific cortical features"

**This PDF file includes:**

Figs. S1 to S21
Tables S1 to S7

***Figure S1***

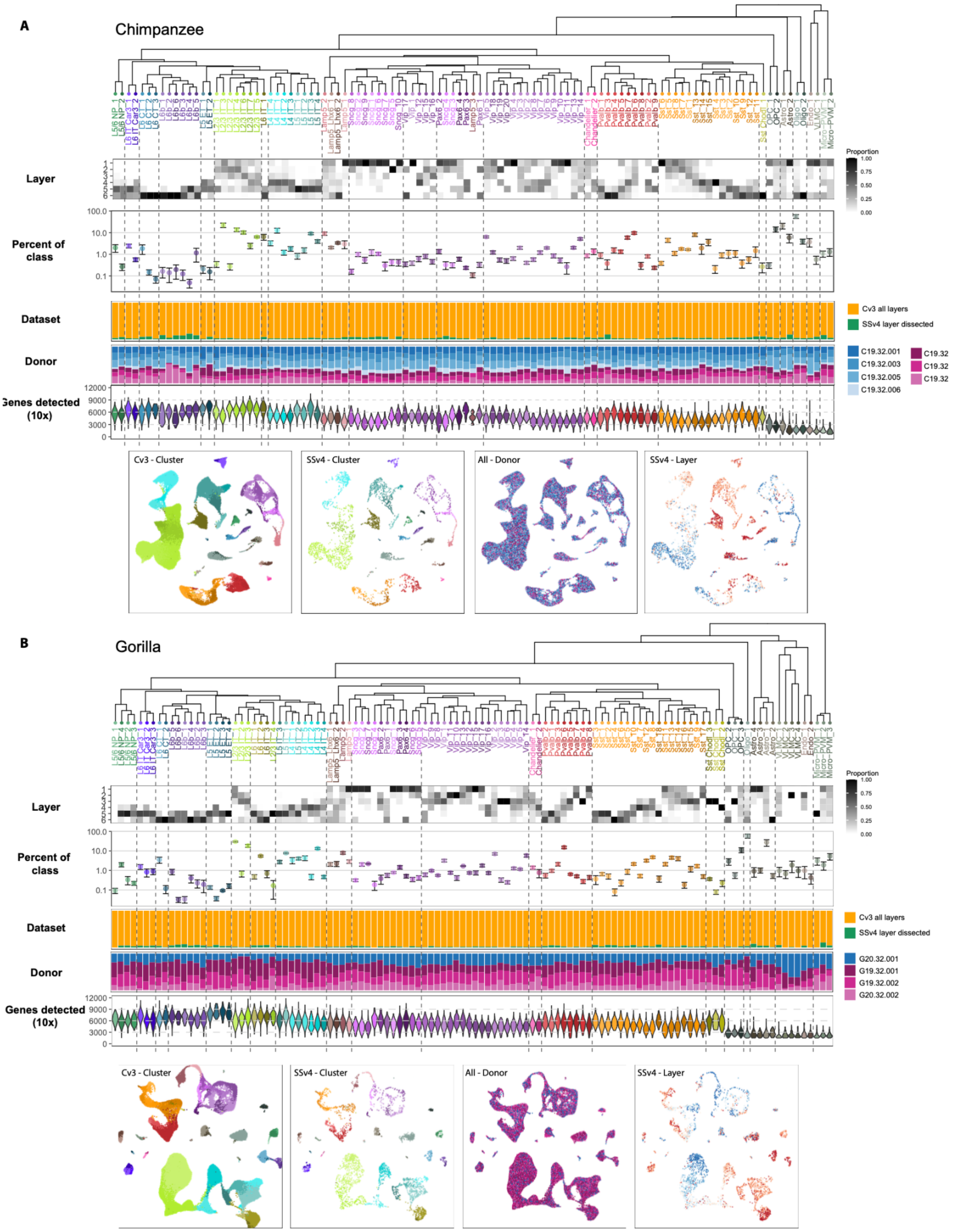


**Fig. S1. Chimpanzee and gorilla MTG taxonomies. (A, B)** Dendrogram of chimpanzee **(A)** and gorilla **(B)** cell types with estimated layer distributions, cell type frequencies as a proportion of cell class, dataset composition, individual ID, and distribution of genes detected. UMAPs of single nuclei colored by cluster, individual ID, and layer dissection.

*Figure S2*

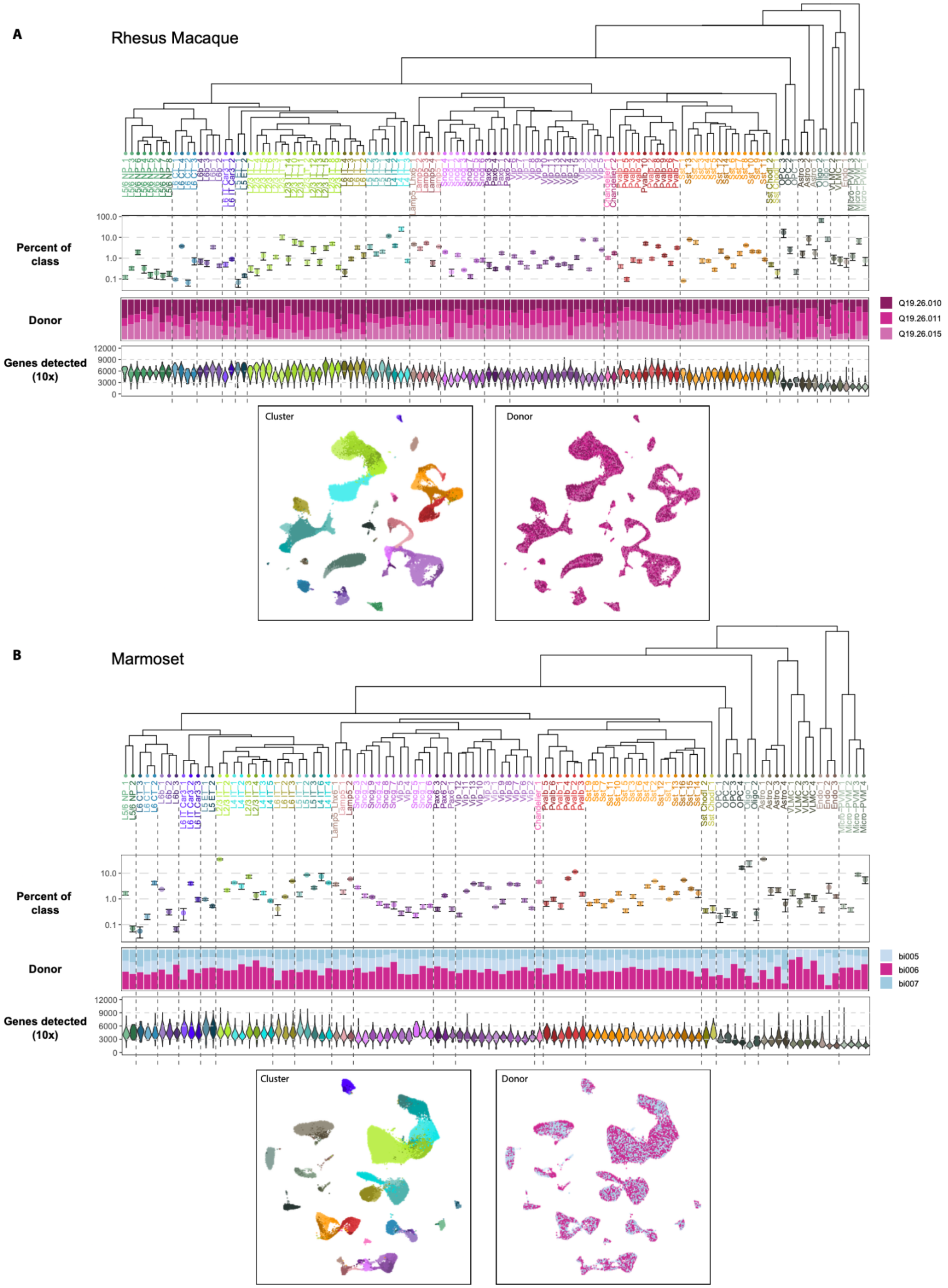
A
Rhesus Macaque
Percent of class
Donor
Genes detected (10x)
Q19.26.010
Q19.26.011
Q19.26.015
Cluster
Donor
B
Marmoset
Percent of class
Donor
Genes detected (10x)
bi005
bi006
bi007
Cluster
Donor

**Fig. S2. Rhesus macaque and marmoset MTG taxonomies. (A, B)** Dendrogram of rhesus macaque **(A)** and marmoset **(B)** cell types with estimated layer distributions, cell type frequencies as a proportion of cell class, dataset composition, individual ID, and distribution of genes detected. UMAPs of single nuclei colored by cluster, individual ID, and layer dissection.

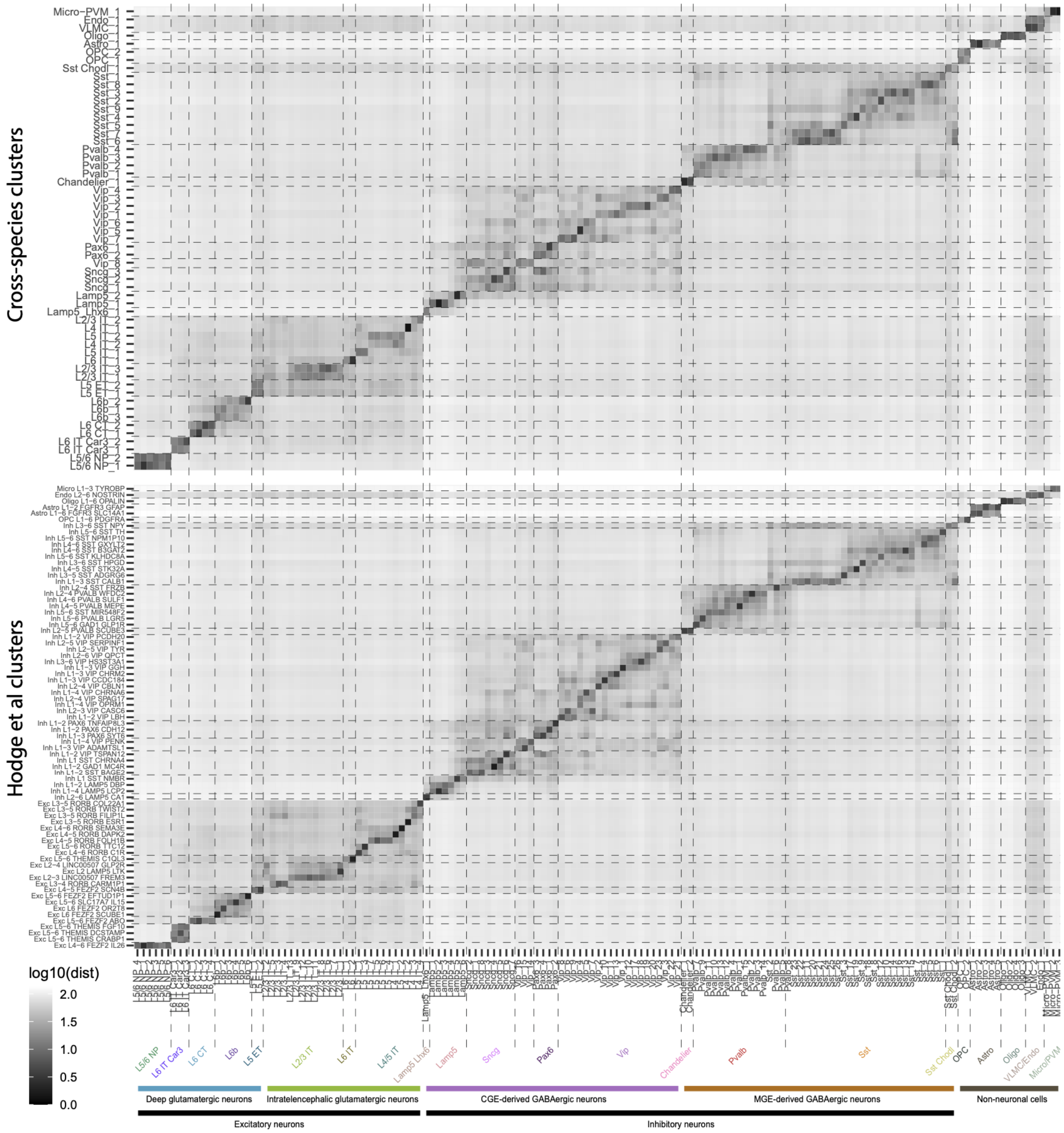


**Fig. S3. Comparison of human MTG cell types to other taxonomies.** Heatmaps of log-transformed Euclidean distance between cluster centroids of human MTG cell types (columns) and primate consensus types (top) and our previously published MTG taxonomy (bottom; Hodget et al. 2019 *Nature*). Cell subclasses, neighborhoods, and classes are labeled at the bottom.

***Figure S4***

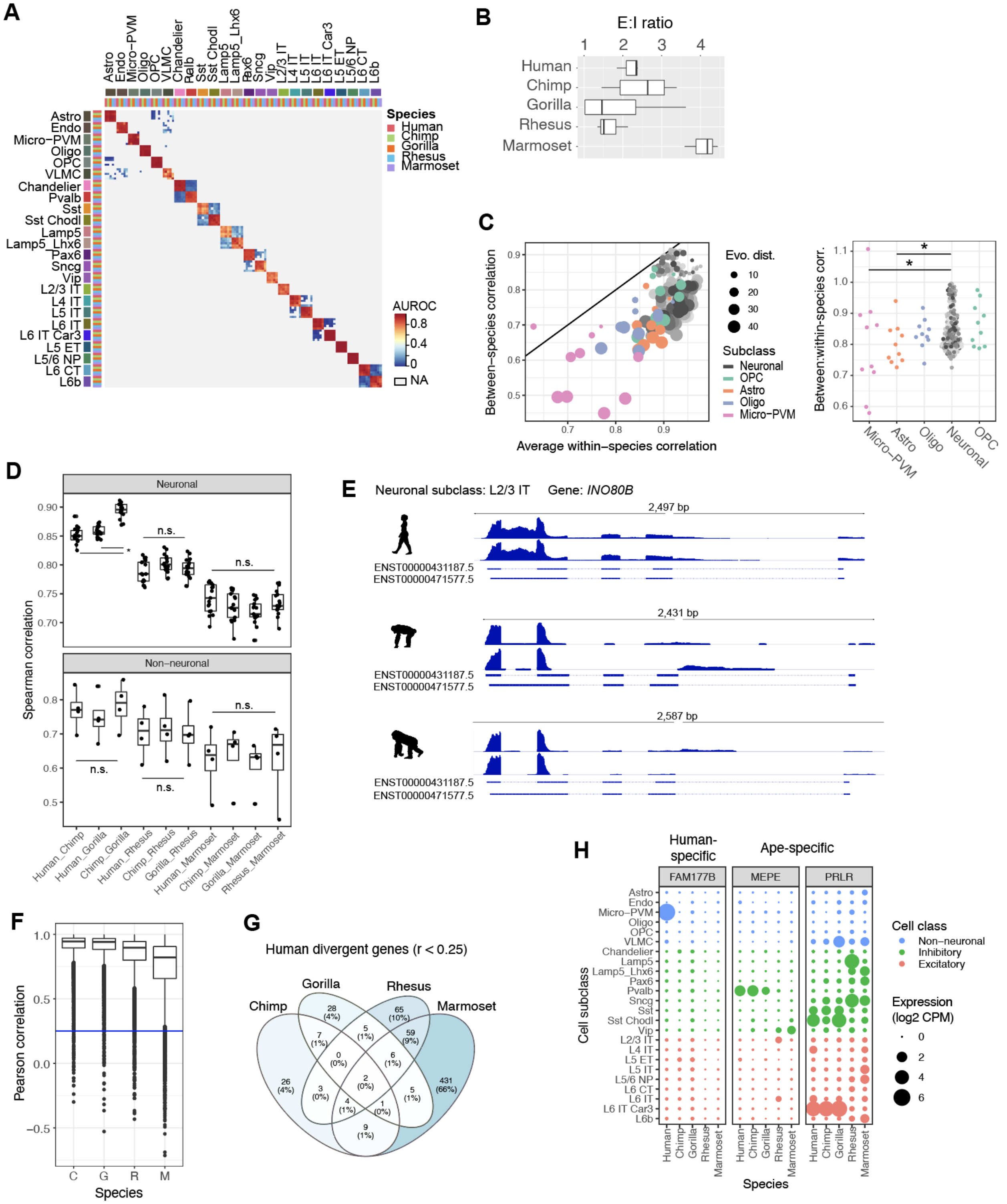


**Fig. S4. Cell subclass comparisons across species. (A)** Heatmap of 1-vs-best MetaNeighbor scores for cell subclasses. Each column shows the performance for a single training group across the three test datasets. AUROCs are computed between the two closest neighbors in the test dataset, where the closer neighbor will have the higher score, and all others are shown in gray (NA). Dark red 3x3 blocks along the diagonal indicate high transcriptomic similarity across all five species. **(B)** Boxplots of the estimated ratio of excitatory to inhibitory neurons across individuals of each species based on snRNA-seq sampling. **(C)** Left: Comparison of cell subclass similarity (expression correlation) within and between species. Each point represents a pairwise species comparison where the point color indicates subclass and size indicates the evolutionary distance to the most recent common ancestor. Within-species similarity was calculated as the average of the average interindividual correlations for the two species being compared. Right: Bee swarm plots showing the ratio of between-species to average within-species correlations for all pairs of species and grouped by non-neuronal subclasses and neurons. The correlation ratio of non-neuronal subclasses was compared to the neuronal ratio using one-sided t tests with Holm-Bonferroni p-value correction for multiple testing (* $P < 0.05$). **(D)** Comparison of expression correlations of neuronal and non-neuronal subclasses across pairs of species. ANOVA tests were used to test for significant differences in correlations across great apes, between great apes and rhesus, and between great apes and marmoset. A post-hoc Tukey HSD test between pairs of great apes showed that chimpanzee and gorilla were significantly more highly correlated than human and chimpanzee or human and gorilla (* $P < 0.05$). **(E)** Smart-seq v4 RNA-seq read pile-ups at the *INO80B* gene locus for L2/3 IT excitatory neurons in human, chimp, and gorilla. Read pile-ups are consistent with human neurons expressing both isoforms, while chimpanzee and gorilla neurons almost exclusively express the second isoform. This is consistent with the isoform proportions estimated using RSEM (see Methods) and visualized in Figure 2H. **(F)** A comparison of gene expression across cell subclasses between human and four NHP species. Boxplots (median, interquartile range - IQR, whiskers, +/- 1.5 IQR, and outliers) summarize the correlations for all one-to-one orthologous genes, and median correlation decreases with evolutionary distance from human. **(G)** Venn diagram of genes with subclass expression correlation < 0.25 between human and one or more species. **(H)** Expression dot plot of example genes with human- or ape-specific expression patterns.

*Figure S5*

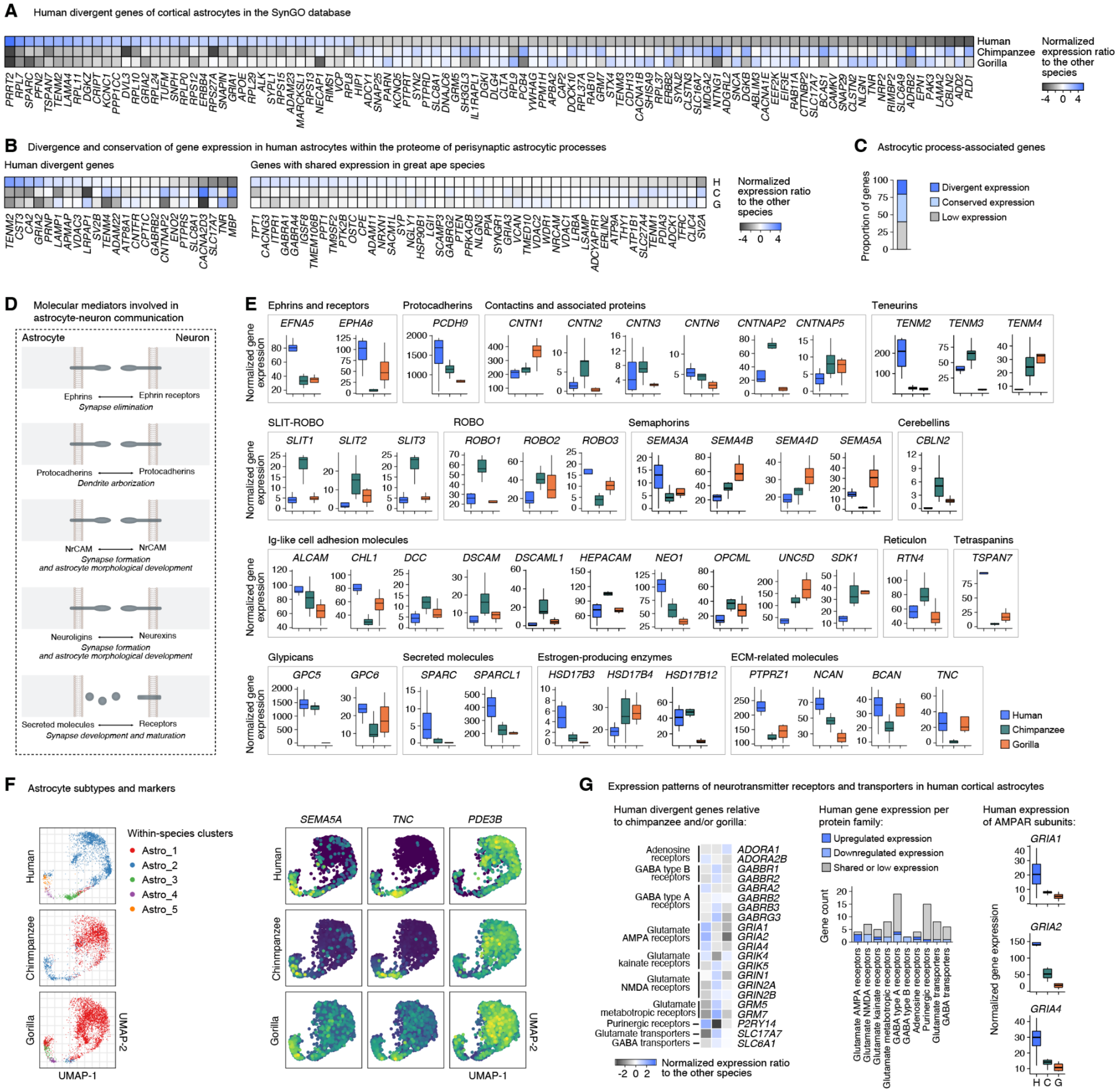


**Fig. S5. Divergent gene programs in astrocytes across great ape species. (A)** Heatmap showing astrocyte SynGO DEGs from the pairwise comparison between human and chimpanzee and the pairwise comparison between human and gorilla (FDR < 0.01, top 100 DEGs visualized). Gene expression change is calculated as the log2 ratio of normalized expression counts in a given species relative to the other two species. **(B)** Heatmap showing genes of the proteome of perisynaptic astrocytic processes identified in Takano et al., 2020 that have divergent expression (left) or conserved expression (right) in human astrocytes compared to chimpanzee and/or gorilla astrocytes (FDR < 0.01). **(C)** Proportion of genes enriched in astrocytic processes from Takano et al., 2020 that show divergent, conserved or low expression in human astrocytes. **(D)** Schematics illustrating the trans-cellular interaction of molecular mediators involved in astrocyte-neuron communication (see Tan et al., 2021 for details on specific molecular pathways mediating astrocytic development and function). **(E)** Examples of DEGs (FDR < 0.05) encoding cell-surface molecules involved in cell adhesion, astrocytic-secreted molecules, and extracellular matrix (ECM)-related molecules that show expression changes in cortical astrocytes across great ape species. **(F)** Markers and subtypes of cortical astrocytes across great ape species. **(G)** Divergent and shared expression of neurotransmitter receptors and transporters in cortical astrocytes. Left, heatmap showing the gene expression changes in human astrocytes compared to chimpanzee and gorilla astrocytes; middle, number of receptors or transporters per gene family that show divergent, shared, or low expression across great ape species; right, human-specific up-regulation of AMPA receptor subunit genes in astrocytes.

### Figure S6

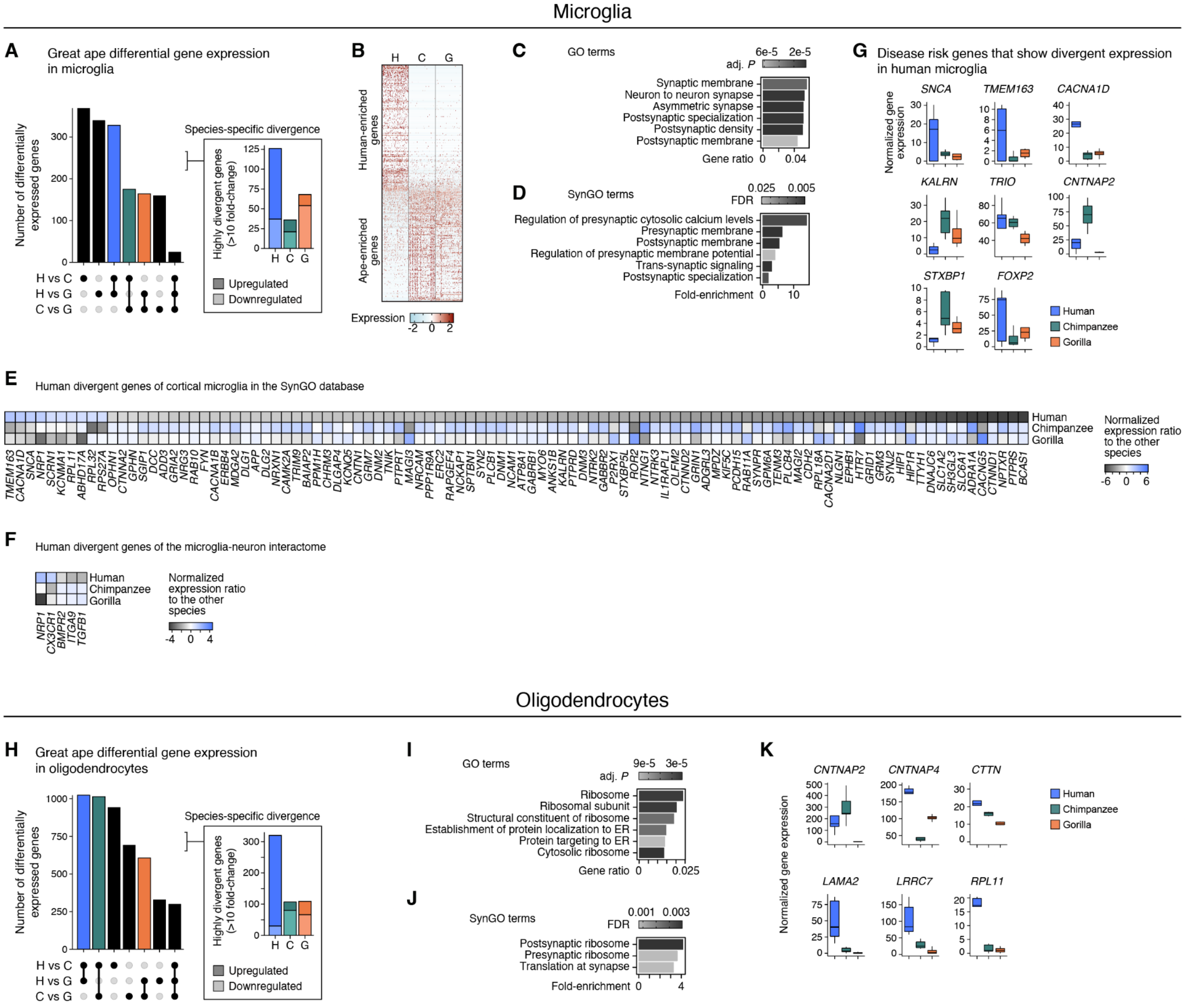


**Fig. S6. Divergent gene programs in microglia and oligodendrocytes across great ape species. (A)** Upset plot showing microglia DEGs for pairwise comparisons of great ape species. **(B)** Heatmap showing row-scaled expression of human versus chimpanzee and gorilla microglia DEGs. **(C-D)** Enrichment of select GO terms **(C)** and SynGO terms **(D)** in the union of microglia DEGs from the pairwise comparison between human and chimpanzee and the pairwise comparison between human and gorilla (FDR < 0.05) that highlights genes related to synaptic processes, ribosomal machinery, and protein targeting pathways. **(E)** Heatmap showing microglia SynGO DEGs from the pairwise comparison between human and chimpanzee and the pairwise comparison between human and gorilla (FDR < 0.01; top 100 DEGs visualized). Gene expression change is calculated as the log2 ratio of normalized expression counts in a given species relative to the other two species. **(F)** Heatmap showing microglia DEGs from the union of the pairwise comparison between human and chimpanzee and the pairwise comparison between human and gorilla (FDR < 0.01) that are found in the microglia-neuron interactome (Stogsdill et al., 2022). **(G)** Examples of disease-linked genes that show gene expression changes (FDR < 0.05) in human microglia compared to chimpanzee and gorilla microglia. **(G)** Upset plot showing oligodendrocyte DEGs across great ape species. **(H-I)** Enrichment of select GO **(H)** and SynGO **(I)** terms in the union of oligodendrocyte DEGs from the pairwise comparison between human and chimpanzee and the pairwise comparison between human and gorilla (FDR < 0.05). **(J)** Examples of DEGs (FDR < 0.01) related to myelin-axon interaction, cell adhesion, and translation that have undergone expression changes in great ape species.

*Figure S7*

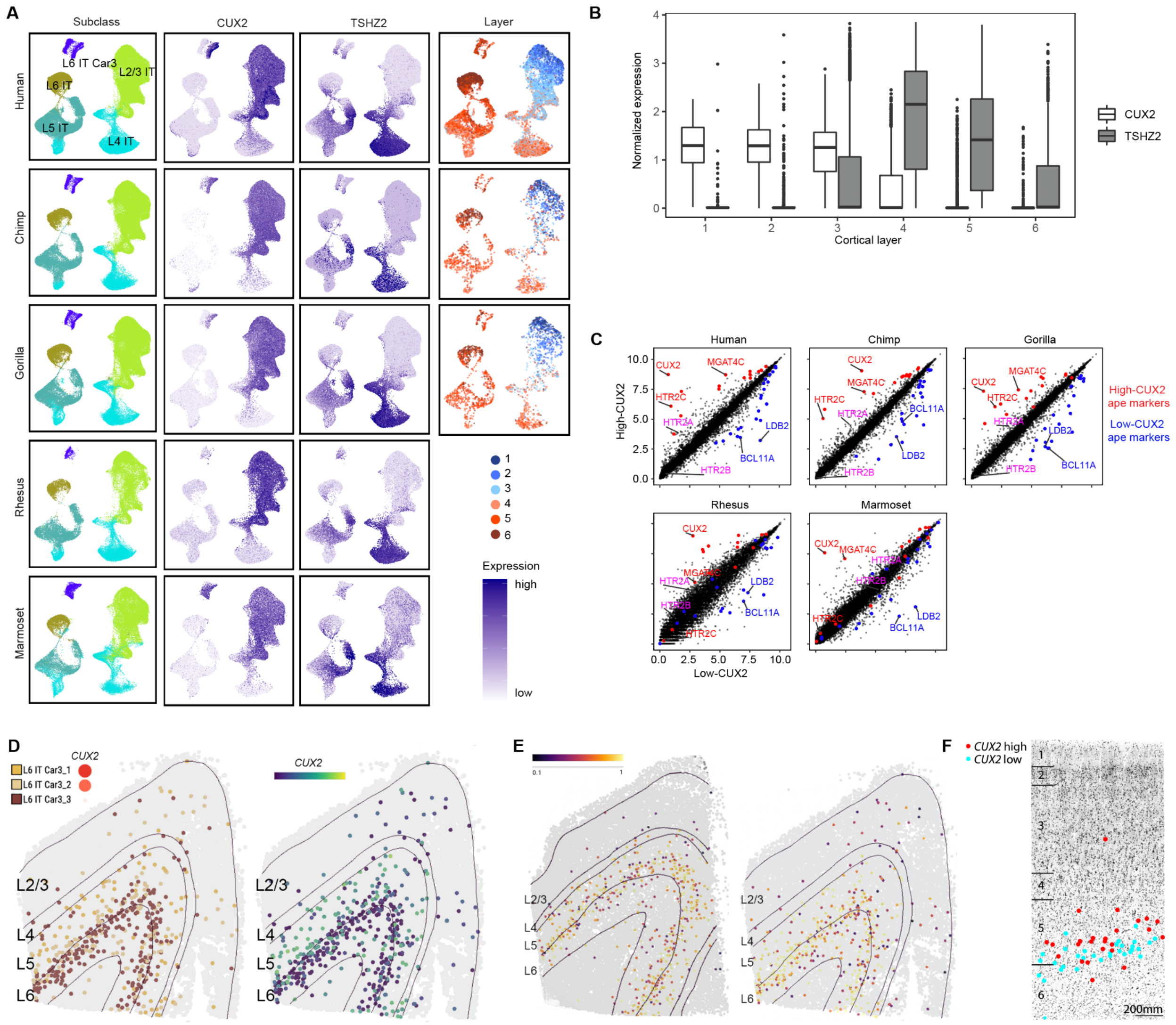


**Fig. S7. Conserved and divergent expression of IT-projecting neurons. (A)** UMAPs of IT-projecting neurons integrated across species and colored by within-species cluster, CUX2 and TSHZ2 expression, and layer dissection. **(B)** Distribution of CUX2 and TSHZ2 expression in human IT neurons dissected from different layers. **(C)** Comparisons of the average expression between High-CUX2 and Low-CUX2 subtypes of L6 IT Car3 neurons in each species. Conserved markers are colored red and blue, and additional serotonin receptor subunit genes are colored pink. **(D)** MERFISH imaging from an additional human individual showing similar proportions and distinct laminar distributions of High-CUX2 and Low-CUX2 subtypes (left) and CUX2 expression in all L6 IT Car3 neurons (right). **(E)** Mapping scores to L6 IT Car3 cell types in the human within-species taxonomy (Fig. 1) measured by marker expression correlations. Tissue sections correspond to the two individuals shown in panel D and Figure 4E. Many neurons in L2/3 have lower mapping scores and may be mislabeled. **(F)** Further in situ validation of laminar distributions of L6 IT Car3 neurons in human MTG using RNAscope to label High-*CUX2* (*CUX2*-high, *LDB2*-, *SMYD1*+) and Low-*CUX2* (*CUX2*-low, *LDB2*+, *SMYD1*+) subtypes based on marker genes. This laminar distribution is consistent with confidently mapped neurons from MERFISH profiling shown in panel E.

*Figure S8*

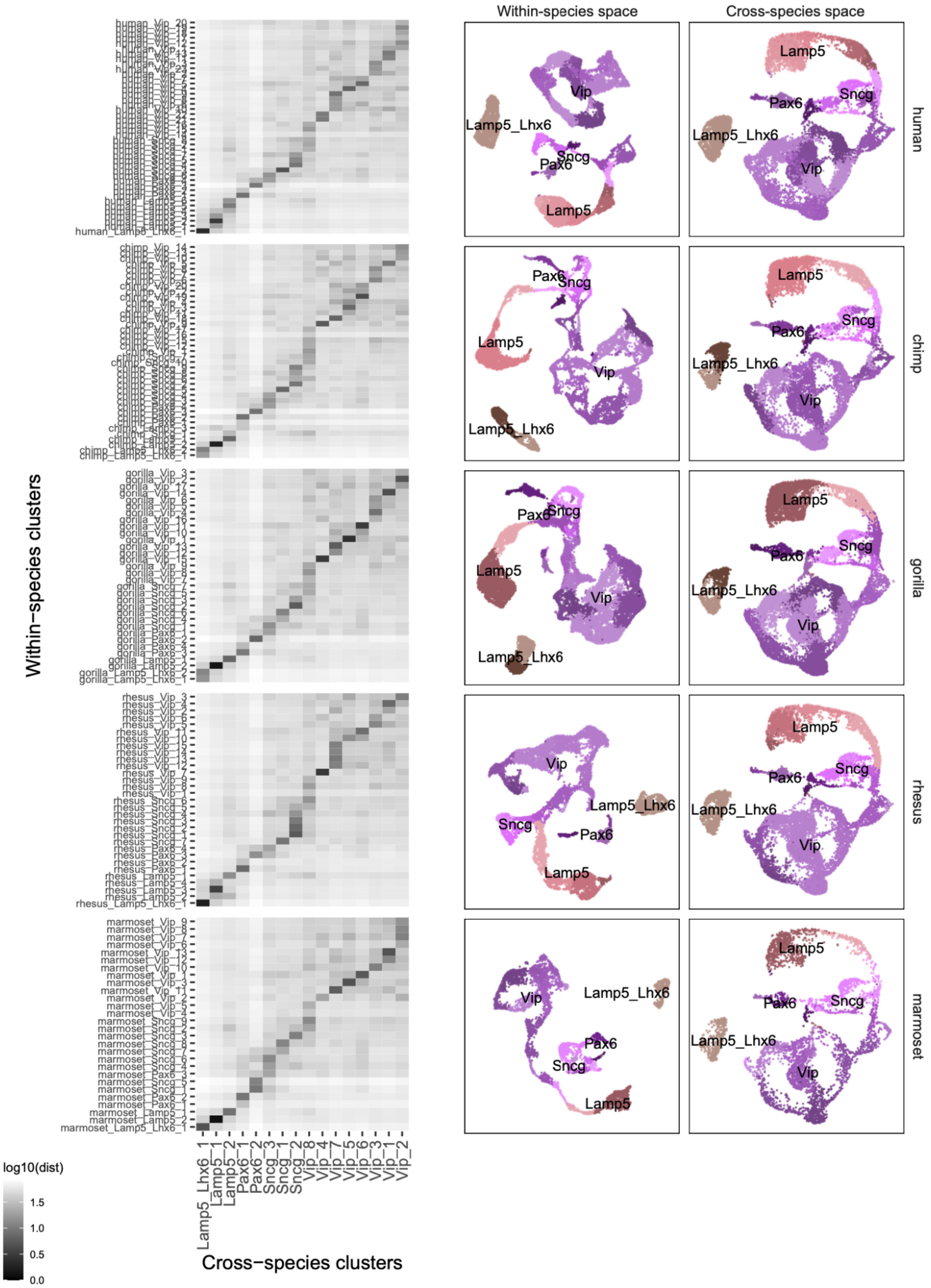


**Fig. S8. CGE-derived neuronal consensus types.** Heatmaps of log-transformed Euclidean distances between within-species cluster centroids and cross-species consensus cluster centroids. UMAPs of single nuclei from each species based on expression within species or integrated across all five species and colored by within species cluster labels.

*Figure S9*

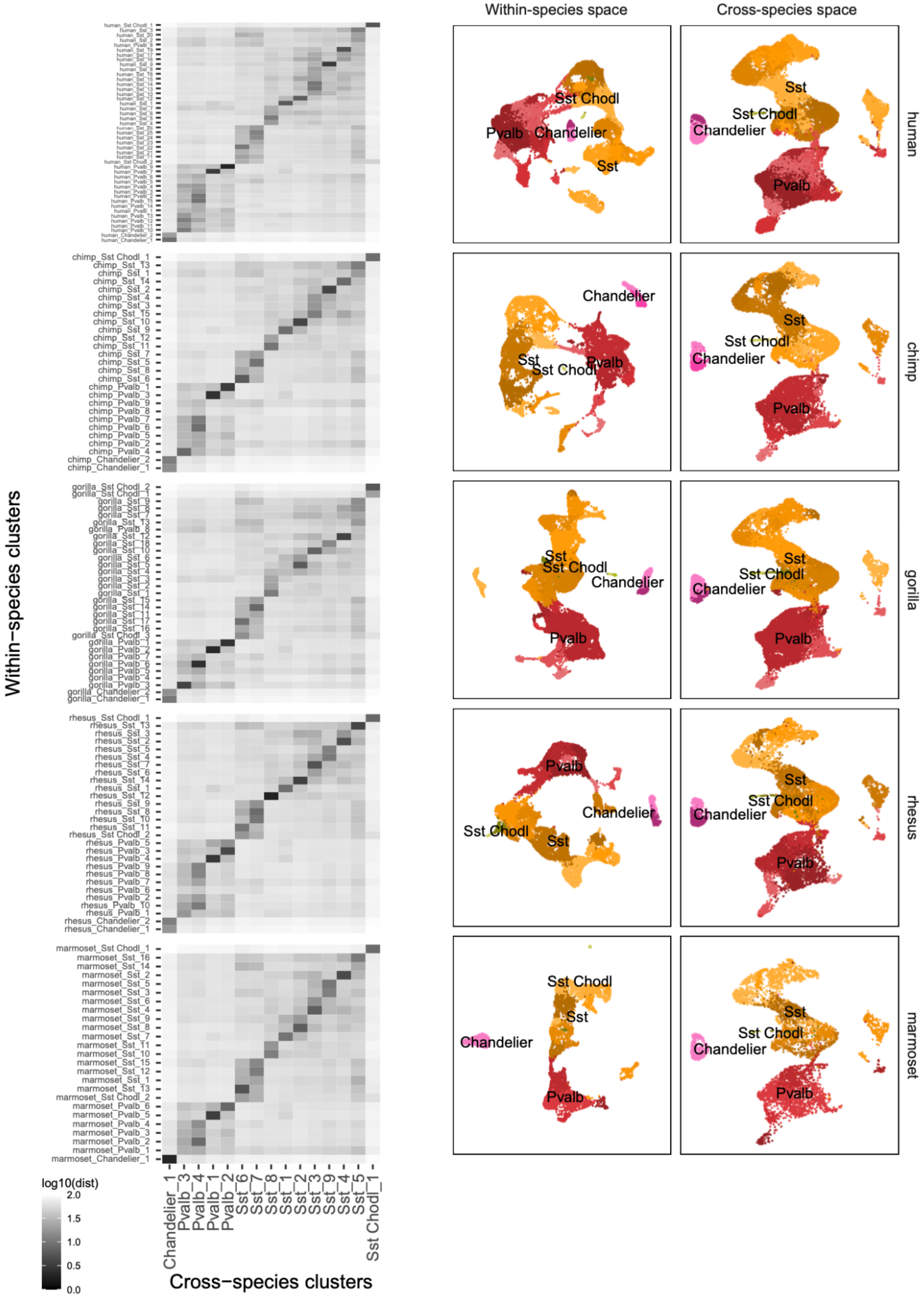


**Fig. S9. MGE-derived neuronal consensus types.** Plots as described for CGE-derived types in **Fig. S8**.

*Figure S10*

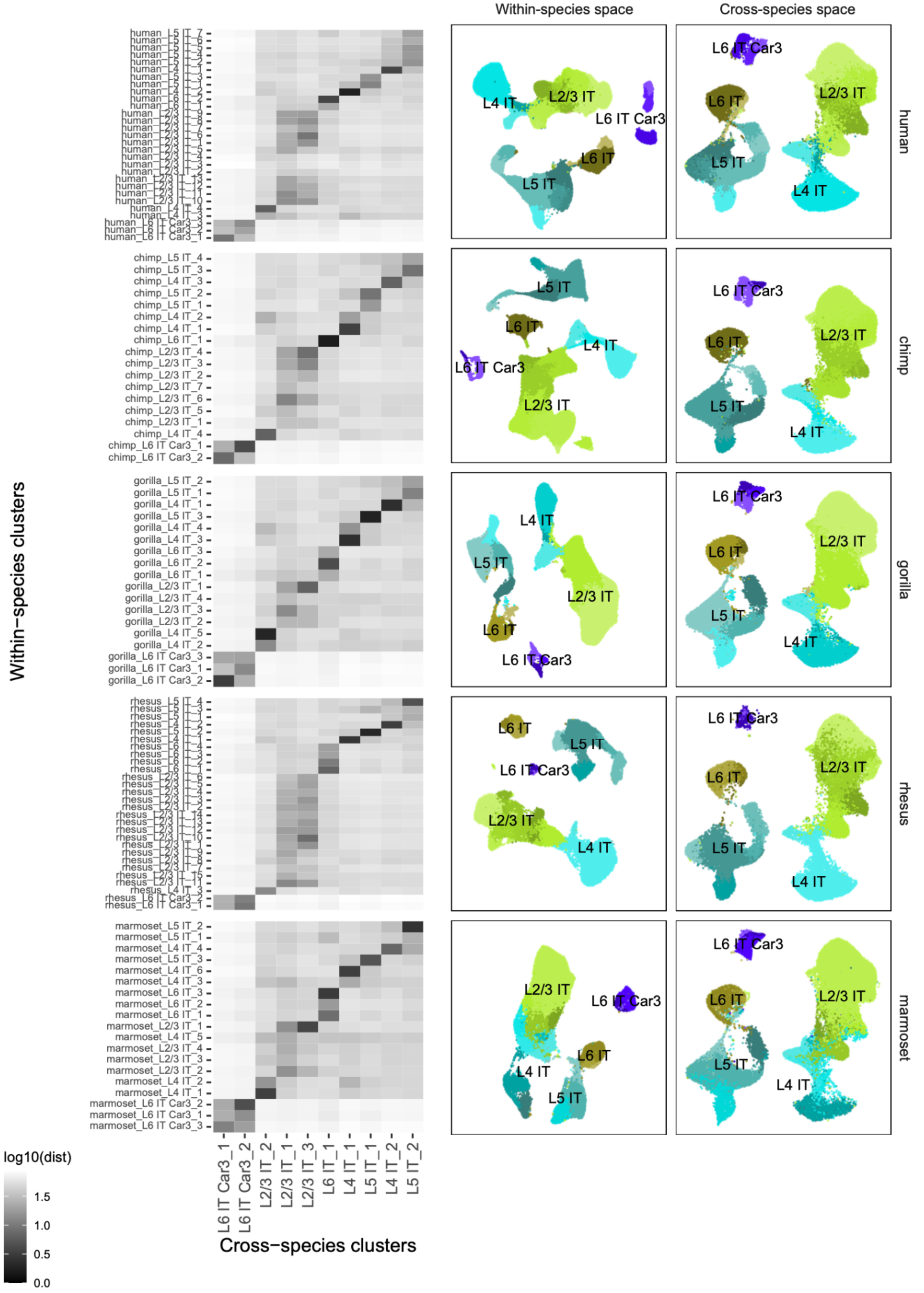


**Fig. S10. Intratelencephalic (IT)-projecting neuronal consensus types.** Plots as described for CGE-derived types in **Fig. S8**.

*Figure S11*

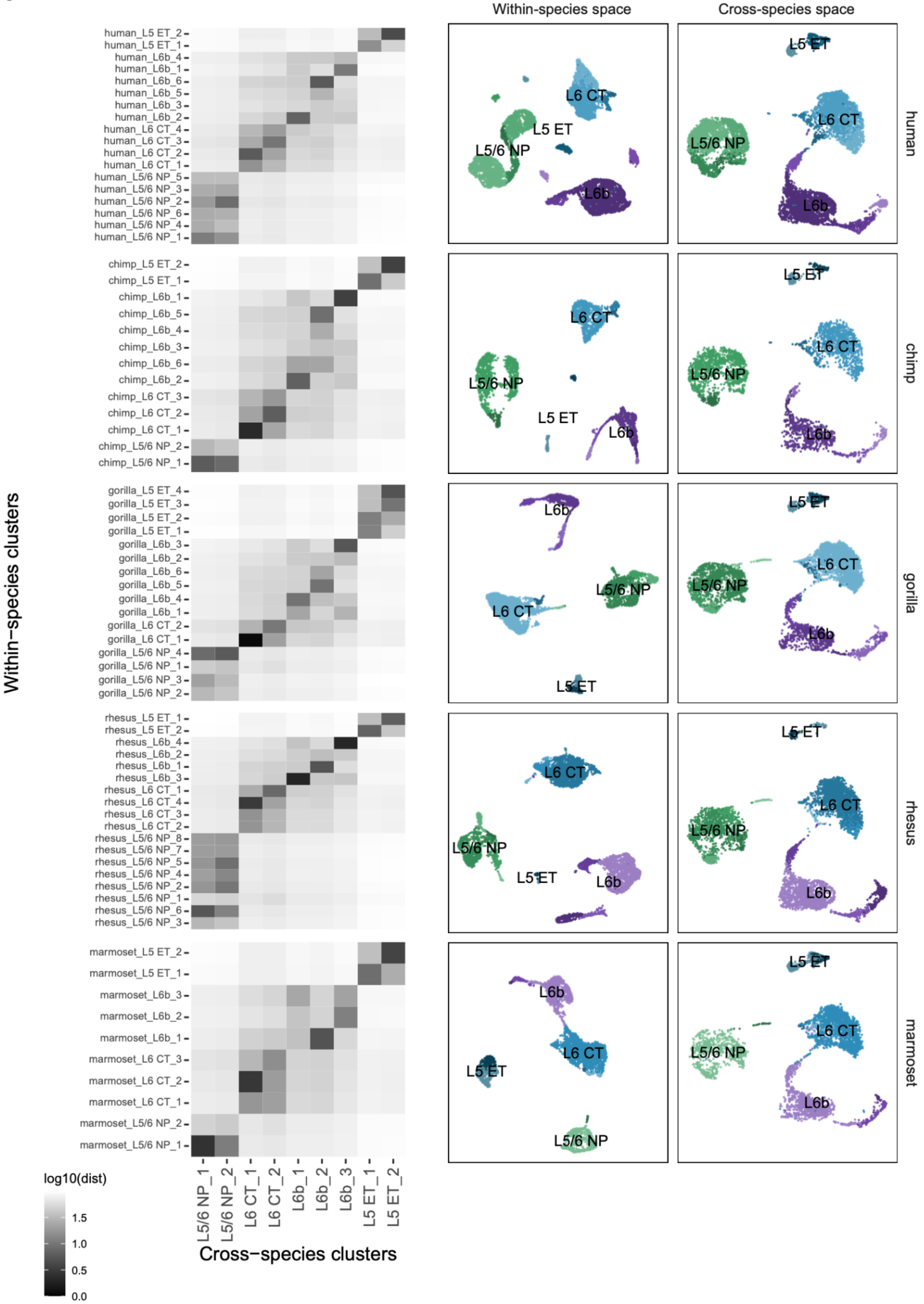


**Fig. S11. Non-IT-projecting neuronal consensus types.** Plots as described for CGE-derived types in **Fig. S8**.

***Figure S12***

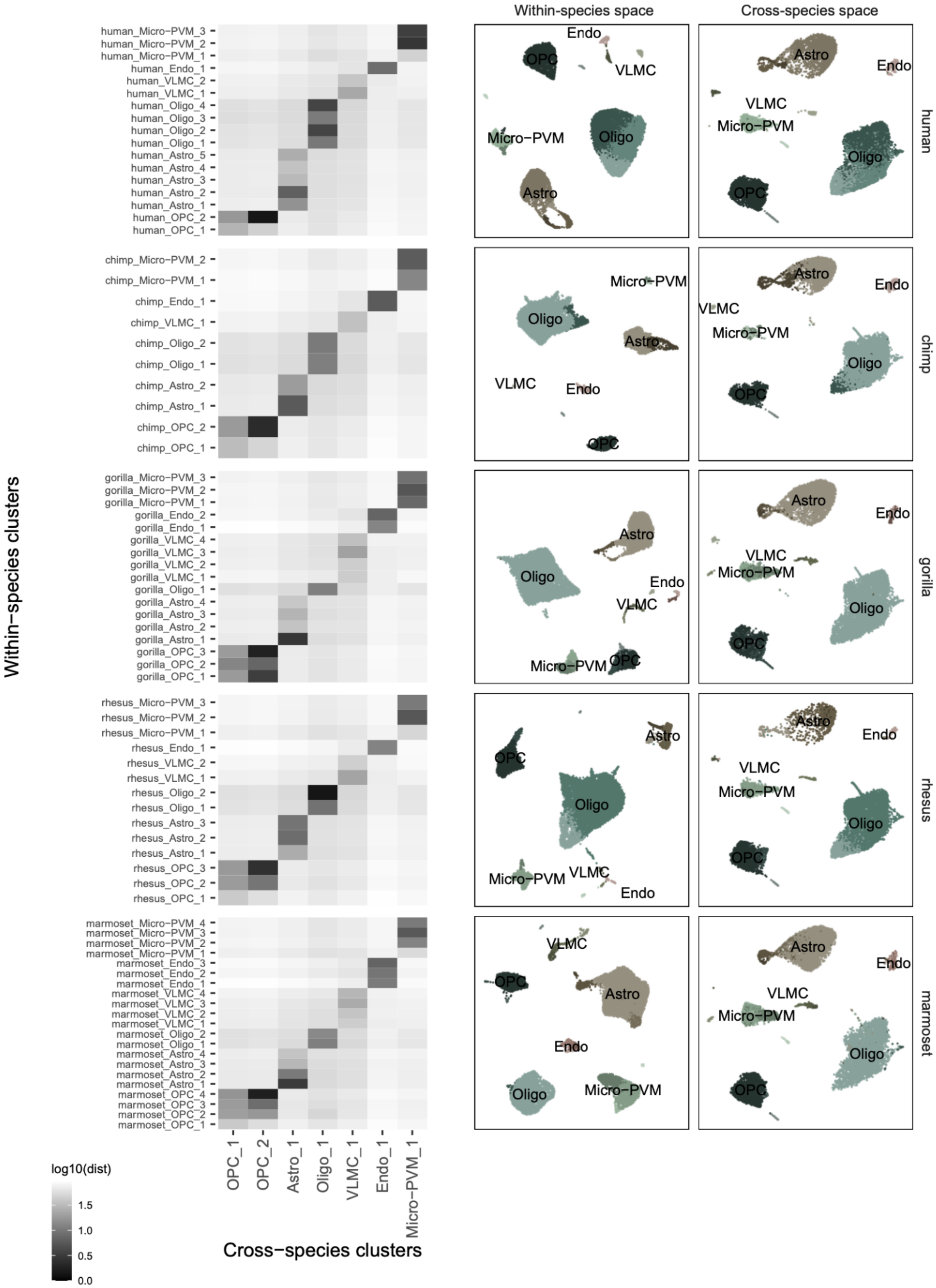


**Fig. S12. Non-neuronal consensus types.** Plots as described for CGE-derived types in **Fig. S8**.

### Figure S13

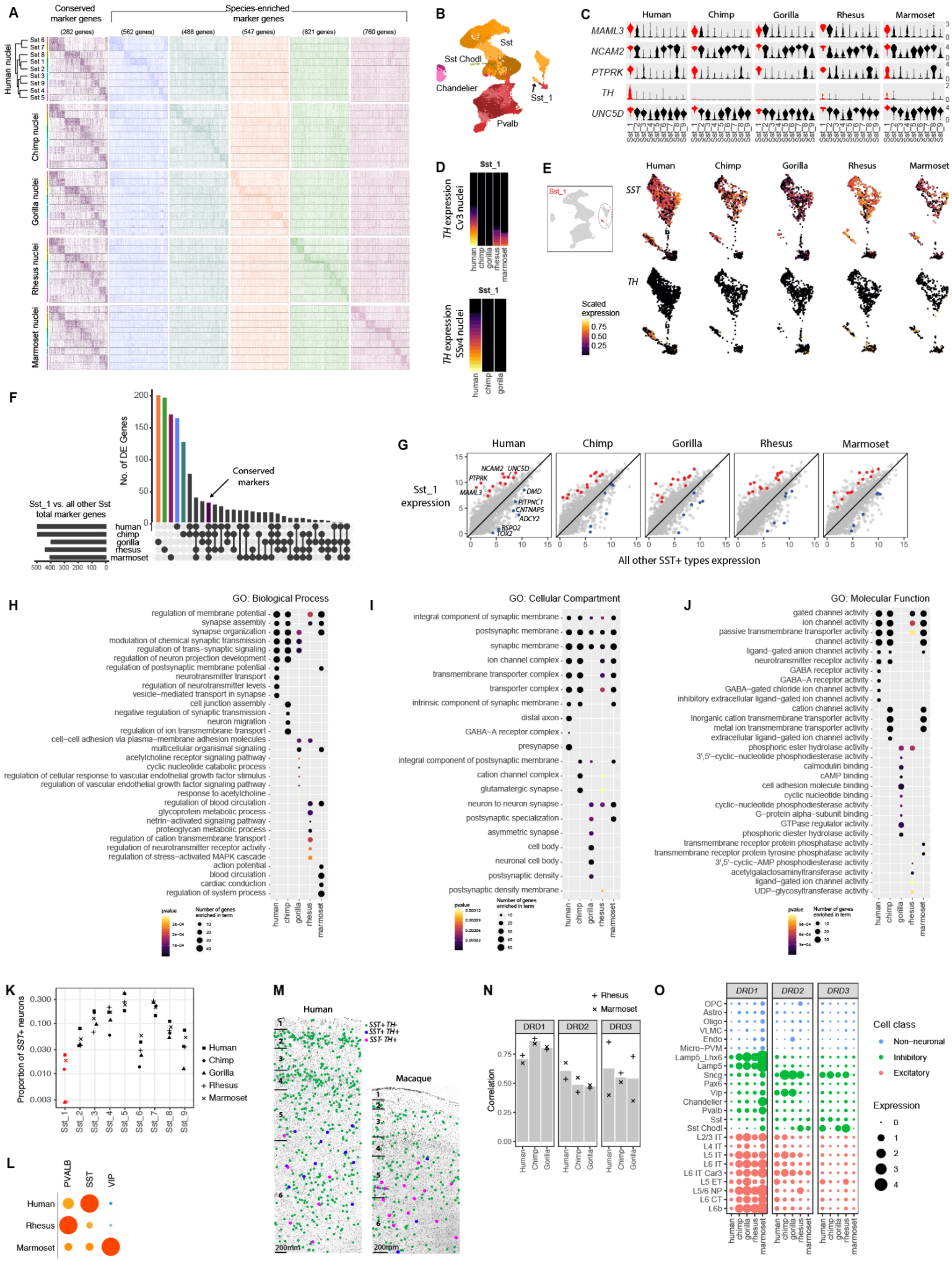

A
Conserved marker genes
Species-enriched marker genes
(282 genes)
(562 genes)
(488 genes)
(547 genes)
(821 genes)
(760 genes)
Human nuclei
Chimp nuclei
Gorilla nuclei
Rhesus nuclei
Marmoset nuclei
B
Sst
Sst Chodl
Chandelier
Sst_1
Pvalb
C
Human
Chimp
Gorilla
Rhesus
Marmoset
MAML3
NCAM2
PTPRK
TH
UNC5D
D
Sst_1
TH expression Cv3 nuclei
TH expression SSv4 nuclei
E
SST
TH
Scaled expression
F
No. of DE Genes
Conserved markers
Sst_1 vs. all other Sst total marker genes
human
chimp
gorilla
rhesus
marmoset
G
Sst_1 expression
All other SST+ types expression
H
GO: Biological Process
I
GO: Cellular Compartment
J
GO: Molecular Function
K
Proportion of SST+ neurons
Human
Chimp
Gorilla
Rhesus
Marmoset
L
PVALB
SST
VIP
M
Human
Macaque
SST+ TH-
SST+ TH+
SST- TH+
N
Rhesus
Marmoset
DRD1
DRD2
DRD3
Correlation
O
Cell class
Non-neuronal
Inhibitory
Excitatory
Expression

**Fig. S13. Conserved SST-expressing interneuron type lacks TH expression in chimpanzees and gorillas. (A)** Conserved and species-enriched markers of SST-expressing consensus interneuron types (**Table S5**). **(B)** UMAP of human MGE-derived interneurons colored by cluster and labeled with major subclasses. Sst_1 labels the SST+/TH+ type. **(C)** Distribution of expression values for marker genes of the Sst_1 cluster (red) versus all other SST-expressing types. **(D)** Stacked bar plots of TH expression measured by droplet-based 3'-UTR sequencing (Cv3) and full length transcript sequencing (SSv4) show no expression detected in chimpanzee and gorilla Sst_1 neurons. **(E)** Inset: UMAP of MGE-derived interneurons integrated across species and colored by Sst_1 cluster membership. Expanded view of SST and TH expression in each species shown for Sst_1 and related types. **(F)** Upset plot summarizing intersections of Sst_1 marker genes across species. A minority of markers are conserved across the five species. **(G)** Average of expression of Sst_1 neurons versus all other SST-expressing types in each species with conserved up-regulated (red) and down-regulated (blue) markers highlighted. **(H-J)** Top enriched GO terms for biological process (H), cellular component (I), and molecular function (J) for species-enriched markers of the Sst_1 type. A minority of terms are significant in all species, and most are species-specific. **(K)** Proportions of SST consensus types among SST-expressing interneurons for each species. Sst_1 neurons are highlighted and range from 0.3–3% of SST+ interneurons. **(L)** Relative proportions (normalized within species) of *PVALB*, *SST*, and *VIP* interneurons that express *TH* in human, macaque, and marmoset. Maximum relative expression is indicated by dot color from low (blue) to high (red). **(M)** *In situ* labeling of cells based on co-expression of *SST* and *TH* in human and macaque MTG using RNAScope. **(N)** Spearman correlations of dopamine receptor subtype (DRD1, DRD2, and DRD3) expression levels across cell subclasses between each great ape species and rhesus and marmoset. Bar plots denote mean correlations for each species and gene, and correlations were not significantly different across species. **(O)** Expression of three dopamine receptor subtypes across cell subclasses and species.

*Figure S14*

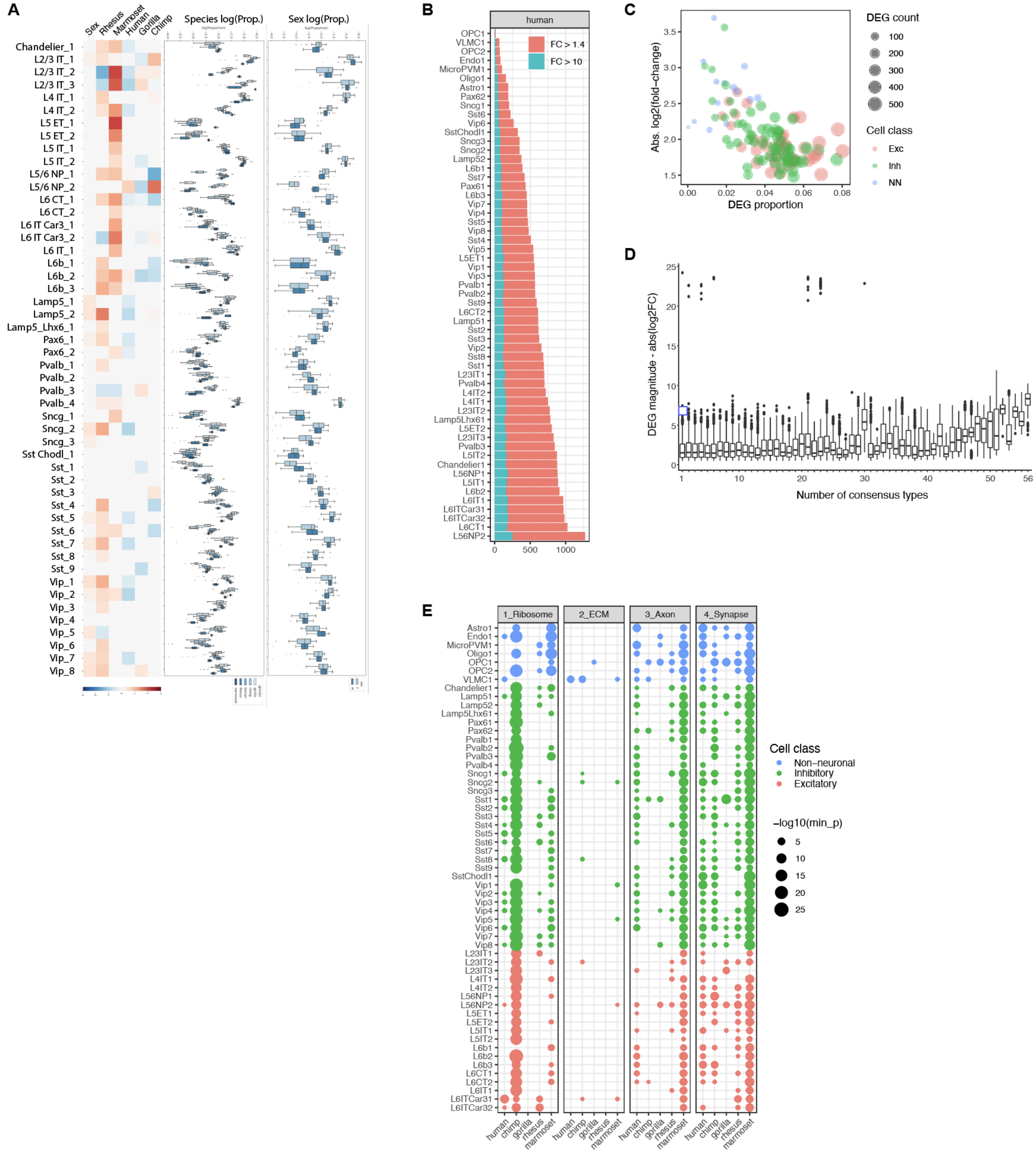

**Fig. S14. Divergent expression of consensus cell types. (A)** scCODA compositional analysis showing neuronal consensus types that change proportions across sex or species. **(B)** Number of hDEGs per consensus type with fold-change (FC) > 1.4 and the subset with FC > 10. **(C)** For consensus type DEGs in humans versus all other species (hDEGs), median log-transformed absolute fold-changes are plotted versus the DEG proportion of median expressed genes in each type. The absolute number of DEGs and cell class are indicated by point size and color, respectively. **(D)** Distribution of absolute fold-change in expression for human DEGs that are grouped by the number of consensus types that show differential expression for those DEGs. Genes that are DEGs in most or all cell types have larger changes in expression. **(E)** Dot plots of consensus types that have significant GO enrichment of DEGs for each species. Dot size denotes the negative log-transformed minimum nominal p-value across all GO terms associated with the four GO categories (ribosome, ECM, axon, and synapse) from **Figure 3**.

***Figure S15***

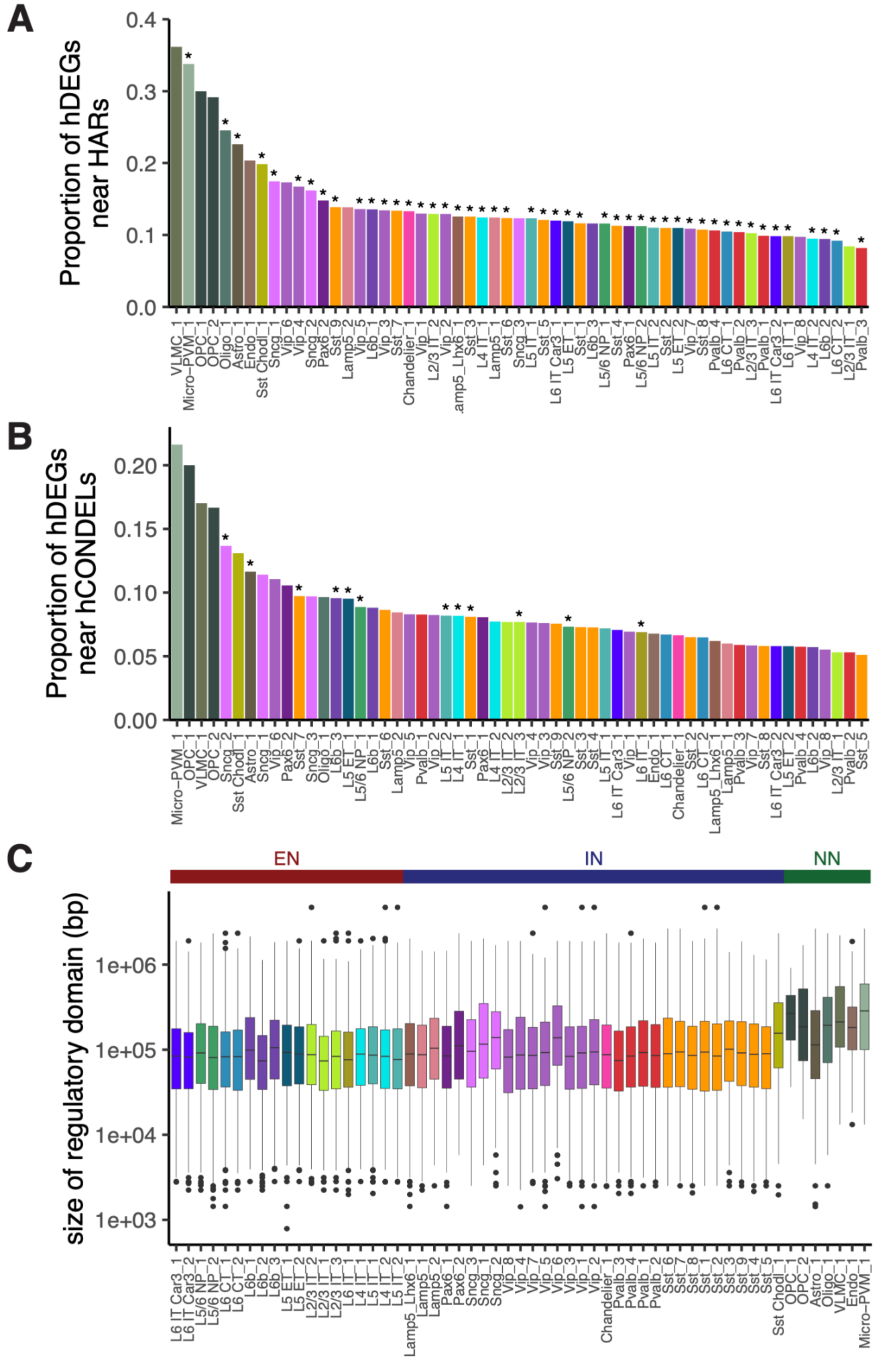


**Fig. S15. Non-neuronal hDEGs have larger gene regulatory domains that contribute to increased enrichment near HARs and hCONDELs.** The regulatory domain for each gene is defined as its intronic regions and its flanking intergenic regions. HARs or hCONDELs within the regulatory domain of a given gene are assigned to that gene. HARs **(A)** and hCONDELs **(B)** are enriched near hDEGs in many consensus types. Asterisks indicate significance at 5% FDR as assessed by the binomial test to account for differences in the size of regulatory domains (*80*). **(C)** Box plots (median and interquartile range (IQR), outliers exceed 1.5 * IQR) of the size distribution of regulatory domains for hDEGs in each consensus type.

***Figure S16***

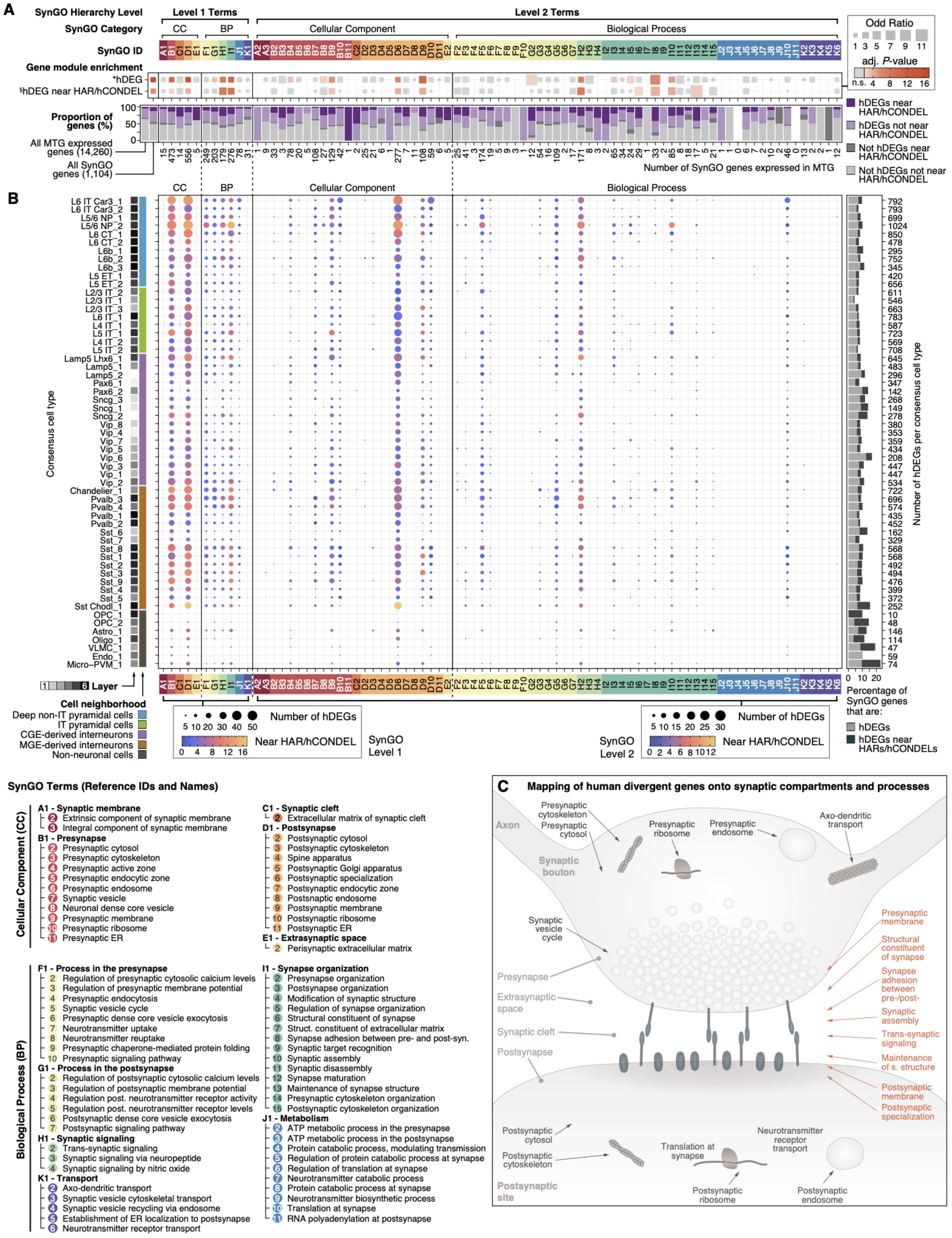


**Fig. S16. Expanded summary of hDEG enrichment near HARs or hCONDELs in SynGO.** The proportion of genes that are hDEGs and/or near HARs or hCONDELs is plotted for all genes expressed in the MTG, all genes annotated in SynGO (*25*), and genes annotated to specific SynGO terms (Table S7) either aggregated across consensus cell types **(A)** or for each individual consensus cell type **(B)**. *Fisher's exact test of whether hDEGs are enriched for a given category compared to all genes expressed in MTG. [§]Fisher's exact test of whether hDEGs near HARs/hCONDELs are enriched for a given category compared to all hDEGs. **(C)** Schematic highlighting SynGO terms that are enriched for hDEGs near HARs or hCONDELs in red. These enriched SynGO terms are concentrated at the synapse.

*Figure S17*

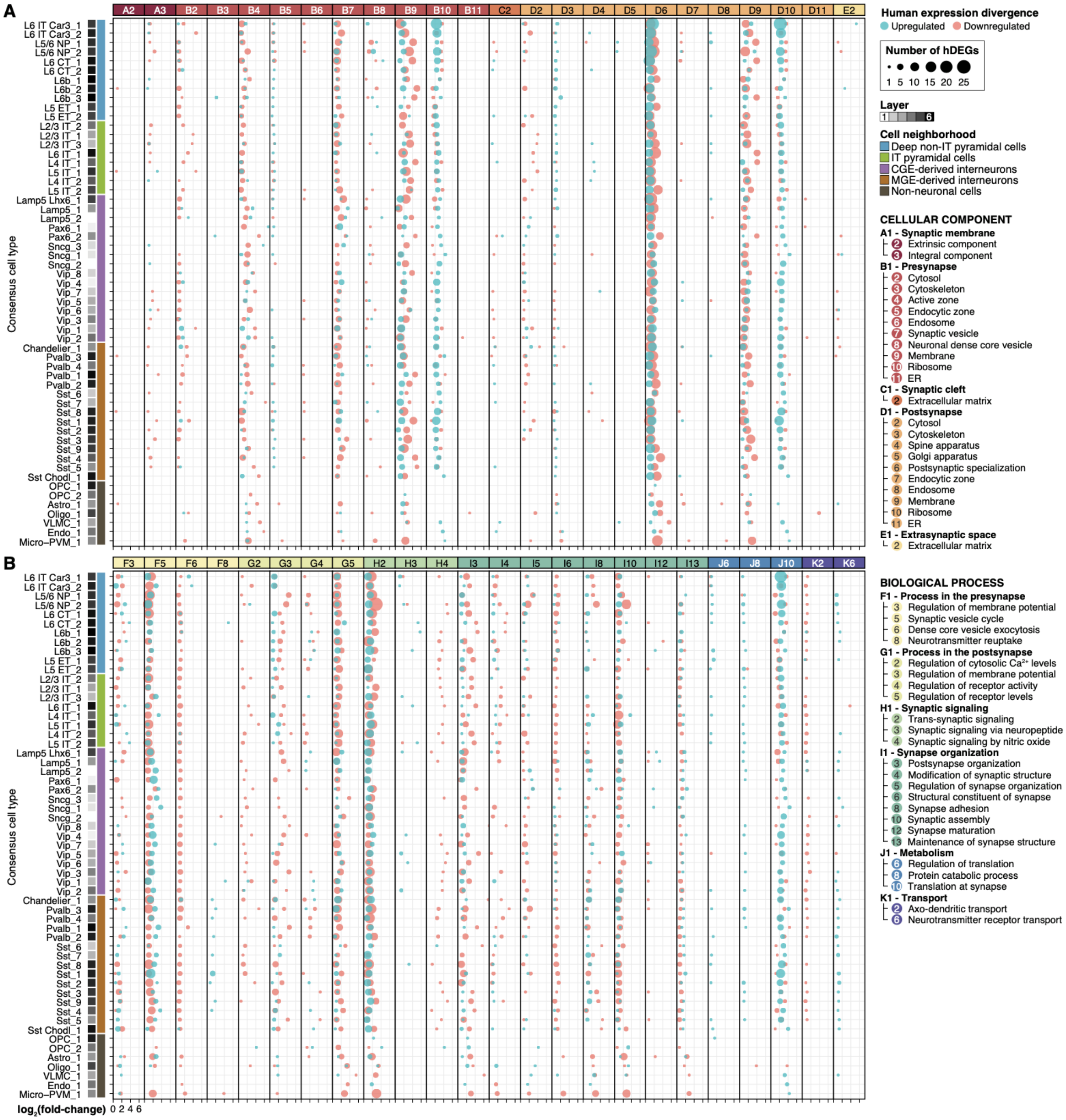


**Fig. S17. Analysis of human gene expression divergence of synaptic compartments and processes across consensus cell types.** Dot plots showing the number of hDEGs in SynGO terms within Cellular Component **(A)** and Biological Process **(B)** categories across consensus cell types. Blue dots represent up-regulated genes, while red dots represent down-regulated genes. Dot sizes indicate the number of hDEGs per consensus cell type and SynGO term.

### *Figure S18*

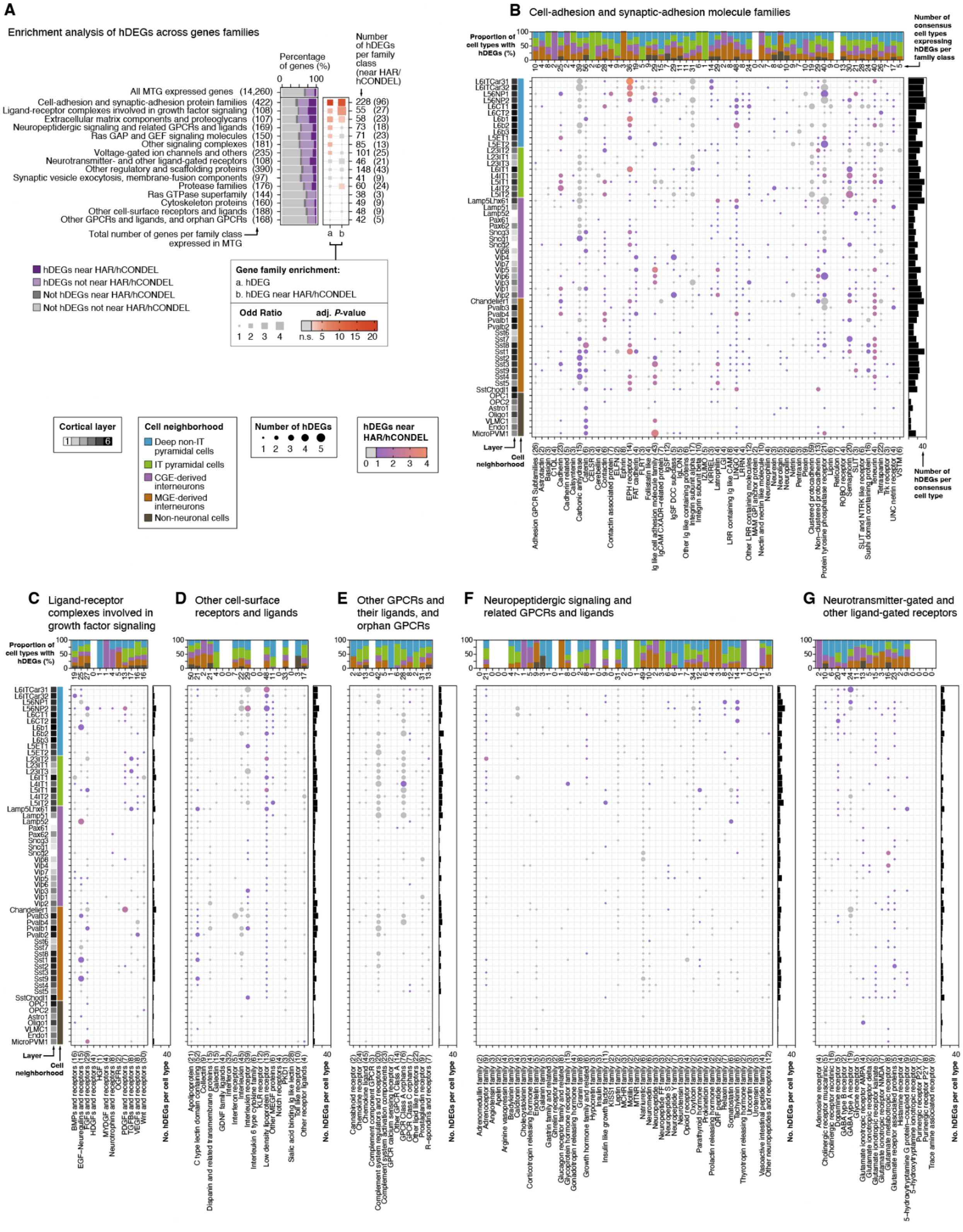


**Fig. S18. Expanded summary of hDEG enrichment near HARs or hCONDELs in gene families. (A)** The proportion of genes that are hDEGs in at least one consensus cell type and/or are near HARs or hCONDELs is plotted for all genes expressed in the MTG and manually-curated lists of gene families aggregated by their likely functional roles at the synapse (Table S7). [a]Fisher's exact test of whether hDEGs are enriched for a given category compared to all genes expressed in MTG. [b]Fisher's exact test of whether hDEGs near HARs/hCONDELs are enriched for a given category compared to all hDEGs. Heatmap colors indicate the significance of category enrichment; gray colors indicate non-significant categories. **(B-G, and continued in Fig. S19)** The number of hDEGs, the number of hDEGs near HARs/hCONDELs, and consensus cell type information is plotted for individual gene families for each category of gene families in (A).

***Figure S19***

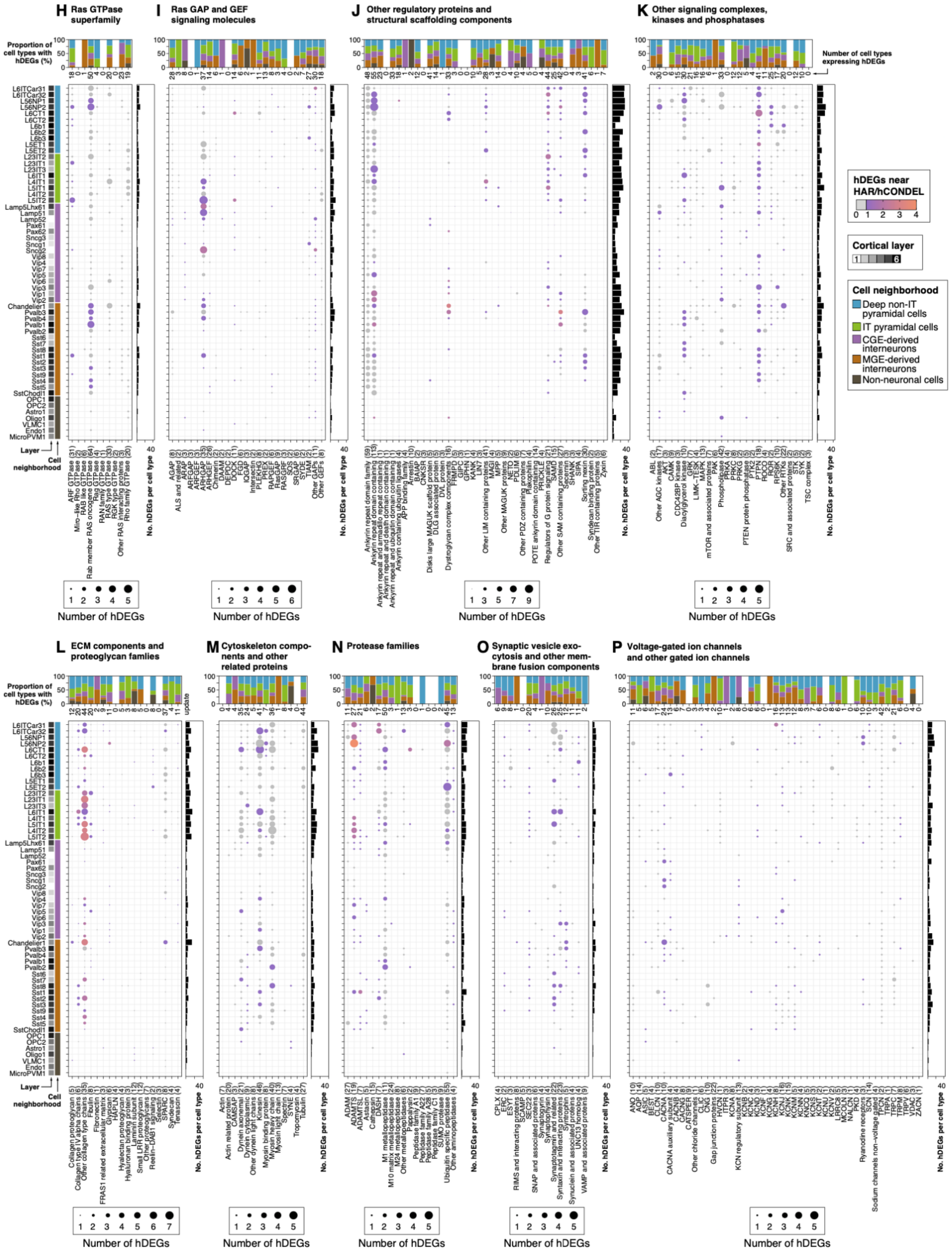


**Fig. S19. Expanded summary of hDEG enrichment near HARs or hCONDELs in gene families (continued).** Continuation of Fig. S18. The number of hDEGs, the number of hDEGs near HARs/hCONDELs, and consensus cell type information is plotted for individual gene families for each category of gene families in Fig. S18A.

### *Figure S20*

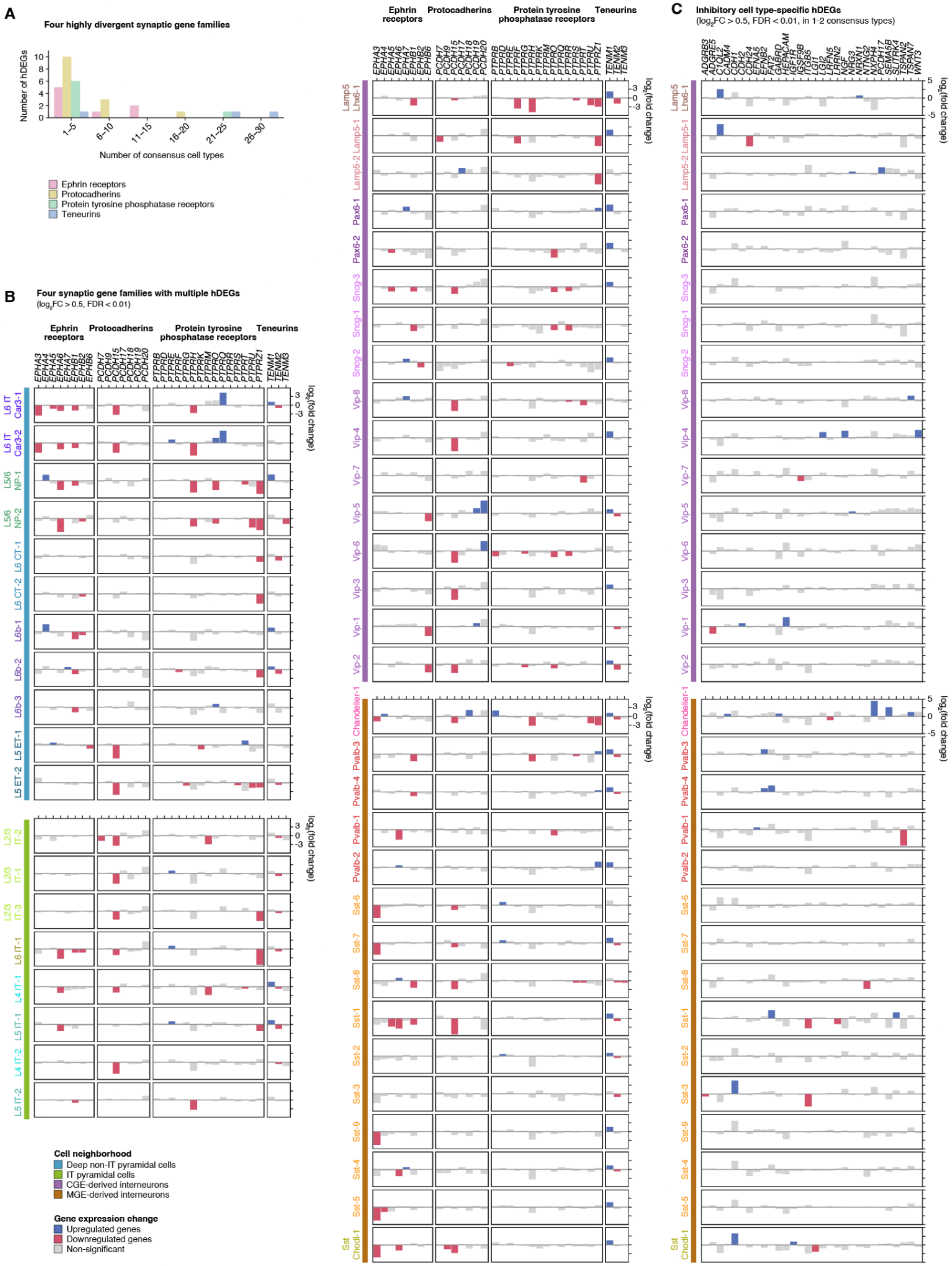
A
Four highly divergent synaptic gene families
Number of hDEGs
Number of consensus cell types
1–5
6–10
11–15
16–20
21–25
26–30
Ephrin receptors
Protocadherins
Protein tyrosine phosphatase receptors
Teneurins
B
Four synaptic gene families with multiple hDEGs
(log2FC > 0.5, FDR < 0.01)
Ephrin receptors
Protocadherins
Protein tyrosine phosphatase receptors
Teneurins
log2(fold change)
L6 IT Car3-1
L6 IT Car3-2
L5/6 NP-1
L5/6 NP-2
L6 CT-1
L6 CT-2
L6b-1
L6b-2
L6b-3
L5 ET-1
L5 ET-2
L2/3 IT-2
L2/3 IT-1
L2/3 IT-3
L6 IT-1
L4 IT-1
L5 IT-1
L4 IT-2
L5 IT-2
Cell neighborhood
Deep non-IT pyramidal cells
IT pyramidal cells
CGE-derived interneurons
MGE-derived interneurons
Gene expression change
Upregulated genes
Downregulated genes
Non-significant
Lamp5 Lhx6-1
Lamp5-1
Lamp5-2
Pax6-1
Pax6-2
Sncg-3
Sncg-1
Sncg-2
Vip-8
Vip-4
Vip-7
Vip-5
Vip-6
Vip-3
Vip-1
Vip-2
Chandelier-1
Pvalb-3
Pvalb-4
Pvalb-1
Pvalb-2
Sst-6
Sst-7
Sst-8
Sst-1
Sst-2
Sst-3
Sst-9
Sst-4
Sst-5
Sst Chodl-1
C
Inhibitory cell type-specific hDEGs
(log2FC > 0.5, FDR < 0.01, in 1-2 consensus types)

**Fig. S20**. **Cell type specificity of human divergent synaptic gene programs. (A)** Number of hDEGs across consensus cell types for four synaptic families that show high degree of divergence in cortical neurons from humans compared to non-human primates. Note that while a few genes (e.g. *TENM2*) are hDEGs in >25 consensus cell types, most members of these families show selective gene expression changes (in 1-5 consensus cell types). **(B)** Gene expression change of members of four highly divergent gene families in human excitatory and inhibitory neuronal types. Y-axis shows scaled fold change of expression in humans relative to non-human primates. Blue indicates human upregulated expression, red indicates human downregulated expression, and grey indicates no significant change in gene expression. **(C)** Examples of synaptic molecules that show gene expression changes specifically in 1-2 inhibitory consensus cell types.

*Figure S21*

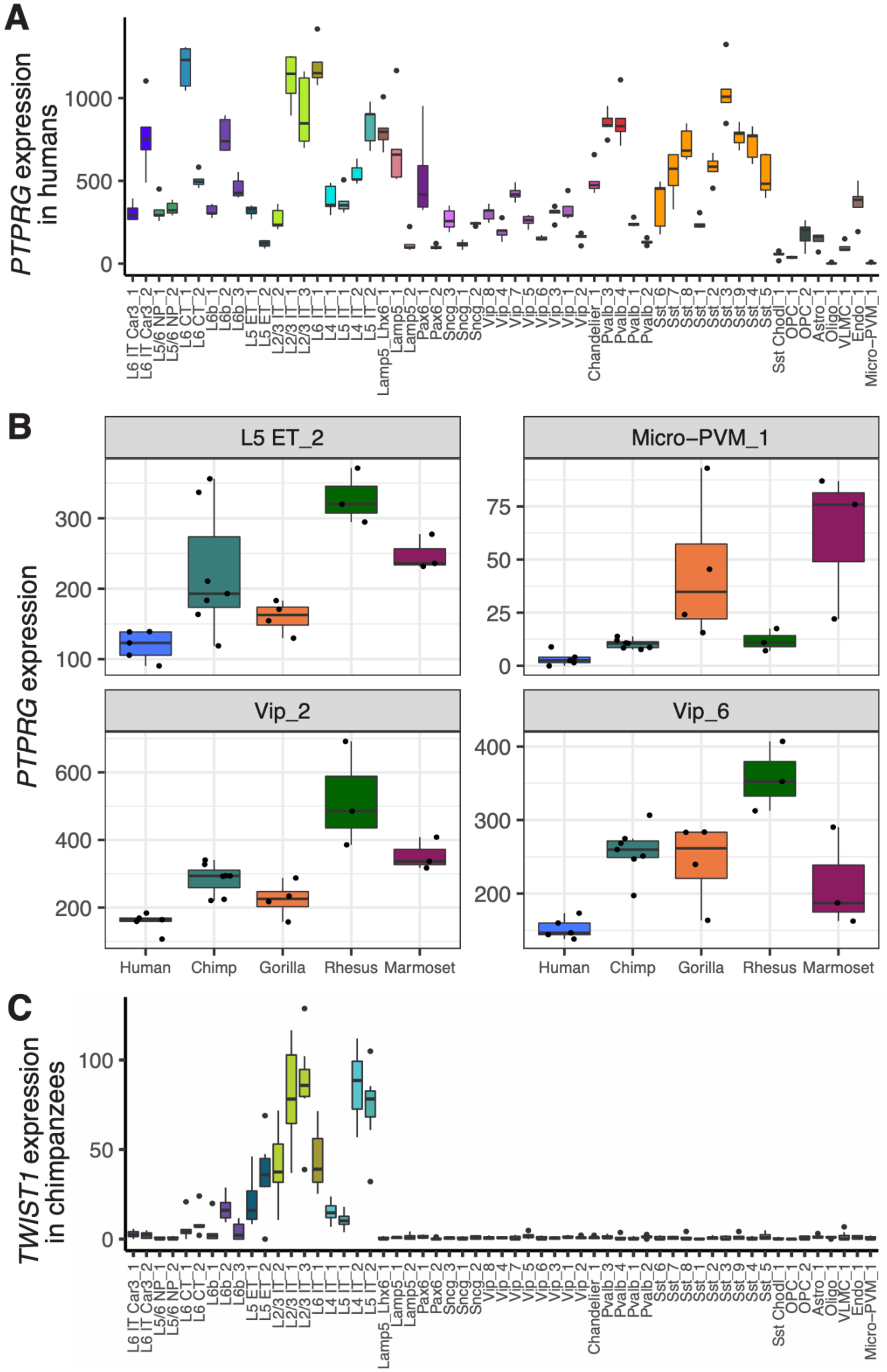


**Fig. S21. *PTPRG* and *TWIST1* expression across consensus types and species. (A)** *PTPRG* expression in each consensus type in humans. **(B)** *PTPRG* expression is decreased in humans compared to NHPs in the consensus types L5 ET_2, Micro-PVM_1, Vip_2, and Vip_6. Points represent normalized *PTPRG* expression per individual. **(C)** *TWIST1* expression in each consensus type in chimpanzees.

***Supplementary Tables***

**Table S1: Within-species cluster proportions across SMARTseq cortical layer dissections.** Estimates of laminar distributions of clusters in human, chimpanzee, and gorilla based on the relative proportions of nuclei dissected from individual cortical layers.

**Table S2: Within-species subclass marker genes.** Prior to DEG analysis, datasets were downsampled to a maximum of 100 nuclei per individual per cluster (finer curation resolution than subclass) in the Cv3 only samples. DEG analysis was performed on log-normalized gene expression matrices. The gene list was generated by implementing Seurat's FindAllMarkers function using the Wilcoxon sum rank test on a maximum of 500 nuclei per group (subclass versus all other cell types) within each species. Genes with Bonferroni adjusted p-values less than 0.05 were retained. This list is not filtered for orthologous genes and reflects the species-specific gene symbol.

**Table S3: Subclass differential gene expression across great apes.** Gene list of pairwise differential expression comparing human, chimpanzee, and gorilla for each subclass. The gene list was generated by implementing a pseudobulk approach across individuals with DESeq2 using Ward's test on downsampled datasets with a maximum 50 nuclei per cluster per individual per species for Cv3 nuclei using one-to-one orthologous genes. Genes with Bonferroni adjusted p-values less than 0.05 were retained.

**Table S4: Subclass isoform switching across great apes.** Relative isoform abundances were compared between human, chimpanzee, and gorilla for one-to-one orthologous genes with at least moderate expression. Gene expression and isoform proportions are reported for two individuals from each species that had SSv4 data. The column 'Large_prop_change' indicates if one species expresses a larger proportion (> 0.7, > 3-fold change on average across individuals) of a transcript than the other species (proportion < 0.1).

**Table S5: SST consensus type markers across species.** List of marker genes that are differentially expressed in each of 9 *SST*-expressing consensus types relative to the other types. Markers were independently generated for each species. The gene list was generated by implementing the Wilcoxon sum rank test with Seurat's FindAllMarkers function on downsampled datasets with a maximum 100 nuclei per cross-species cluster per individual for Cv3 nuclei.

**Table S6: Consensus type species-specific expression.** Gene list of differential gene expression comparing each species to the other four species for each consensus type. The gene list was generated by implementing a pseudobulk approach across individuals with DESeq2 using Ward's test on downsampled datasets with a maximum 50 nuclei per cluster per individual per species for Cv3 nuclei using one-to-one orthologous genes. Genes with Bonferroni adjusted p-values less than 0.05 were retained.

**Table S7. Analysis of hDEGs near HARs or hCONDELs.** Lists of hCONDEL coordinates, hDEGs in consensus types that are near HARs or hCONDELs, potential HAR-hDEG interactions from existing PLAC-Seq datasets, and manually curated synapse-related gene families. This table also includes statistics on the enrichment of hDEGs, and hDEGs near HARs or hCONDELs, for SynGO and for manually curated lists of synapse-related gene families.
